# NeuroDisco: Interfacing neuronal populations - cell by cell, millisecond by millisecond, day by day

**DOI:** 10.64898/2026.11.08.764989

**Authors:** Cyprian S. Adler, Friedrich Schwarz, Hannah Zoé Kerger, Hari Prasad Sreekrishnapurath Variyam, Tabea Ruschmeyer, Laslo Addy Bode, Felix Preuss, Julian Vogel, Viola Prieseman, Markus Schönekeß, Joséphine Barbéry, Laura Diaz, Fred Wolf, Manuel Schottdorf, Andreas Neef

**Affiliations:** 7Göttingen Campus Institute for Dynamics of Biological Networks, Heinrich Düker Weg 12, 37073 Göttingen, Germany; Max Planck Institute for Dynamics and Self-Organization, 37077 Göttingen, Germany; Institute for Neuropathology, University Medical Center Göttingen, 37075 Göttingen, Germany; Bonn-Cologne Graduate School for Physics and Astronomy (BCGS), University of Cologne; III. Institute of Physics, University of Göttingen, 37077 Göttingen, Germany; Max Planck Institute for Multidisciplinary Sciences, 37075 Göttingen, Germany

**Author notes:** These authors contributed equally to this work and share first authorship (Cyprian S. Adler, Friedrich Schwarz). These authors contributed equally to this work and share senior authorship (Manuel Schottdorf, Andreas Neef).

## Abstract

Neuronal cultures offer comprehensive access to living networks, but their value as functional models is limited by synchronized bursts that drive pathologically strong neuronal depolarization. Theory attributes such bursting to a lack of external input. Here we present NeuroDiscos, compact µLED and OLED projectors that operate inside standard incubators and deliver patterned, long-term optogenetic stimulation, synchronized with submillisecond precision to multi-electrode array recordings. Weak, uncorrelated NeuroDisco stimulation dispersed the bursts but left the population firing rate largely unchanged. In this asynchronous regime, neuronal activity reproducibly follows the weak stimulation, and stimulus-response relations characterize receptive fields and putative functional connections. At hardware costs of €500-1,000, NeuroDiscos offer a scalable route to functional characterization of primary and patient-derived cultures.

## Introduction

Neural circuits operate on timescales ranging from the millisecond precision of spike timing to the hours over which synapses and excitability adapt. Neuronal cultures have been instrumental in uncovering such mechanisms, including spike-timing-dependent plasticity ^1^, homeostatic plasticity ^2^, and activity-dependent synaptic remodeling ^3^. Compared with the intact brain, neuronal cultures offer greater experimental accessibility and, in principle, the possibility of combining precisely defined external inputs with continuous recordings from large parts of a network over days. Patient-derived networks now extend this access to human disease modeling and drug testing ^4,5^.

In practice, however, culture activity is dominated by synchronized population bursts that recruit most of the network ^6,7^. Although these bursts resemble epileptiform discharges caused by massive synaptic input ^8^ and are largely absent in vivo, their properties are often used as functional readouts of primary and patient-derived cultures ^9,10^. Theory attributes bursting to the near absence of external input in vitro and predicts that weak, sustained input moves networks toward irregular, asynchronous dynamics ^11^. Interventions in neuronal cultures have not achieved a stable burst reduction: distributed electrical stimulation suppresses bursts only during minute-long runs while raising tonic firing ^12^, and neuromodulatory desynchronization gives way to renewed synchrony within hours ^13^.

Ideally, stimulation of neuronal cultures would not only alleviate the problem of synchronous bursting but complement electrical spike detection to create a multi-functional device to condition, interrogate and train neuronal networks. High-density microelectrode arrays record at near single-cell resolution over long periods ^14–17^. A matching stimulator would (1) be synchronized with the recording but not interfere with it, (2) operate stably for days, (3) combine broad spatial coverage with near single-cell resolution and (4) be robust, affordable and compatible with standard culture and recording hardware, so that it can be scaled within standard incubators. Electrical stimulation produces artifacts ^18^, and usually only allows for a limited number of parallel stimulation sites ^16^. In addition, it lacks cell-type specificity, even where it addresses single neurons ^19^. Optogenetic stimulation resolves these limitations, but existing implementations meet only some criteria. Whole-field LED illumination is inexpensive and has been used for hours to days ^20–22^ but lacks spatial structure (criterion 3). Microscope-based pattern illuminators reach single-cell resolution ^23–25^ but are costly, usually operate outside the incubator and have been used only for minutes to about an hour (criteria 2 and 4). Light sources attached to or built into the culture substrate ^26,27^ are compact but integrate poorly with standard culture and recording equipment, which hinders scaling (criterion 4).

Here we present NeuroDiscos, compact projectors that project a µLED or OLED array onto light-sensitized neurons growing on multi-electrode arrays inside a humidified incubator. The flexND projects 16 × 16 µLEDs as ∼90 µm spots through a glass-bottom 60-electrode MEA; the HD-ND projects 2 million pixels of an OLED display through a custom lid onto a 26,400-electrode CMOS array. Frames are synchronized to the recording with submillisecond precision at irradiances of 1-20 µW/mm^2^. The hardware costs are about €500 and €1,000 per unit, and up to eight of the small devices can run in parallel in one incubator for days to weeks.

We first characterize synchronization, optical resolution and long-term compatibility with cultured neurons. We then show that weak, spatiotemporally uncorrelated stimulation disperses network bursts and shifts them off their intrinsic propagation paths while leaving the population firing rate largely unchanged. In the resulting asynchronous regime, we characterize neurons’ spatiotemporal receptive fields and find that they capture dendritic structure and putative functional connections. Although patterned stimulation does not increase the population firing rate, it tightly controls the spiking pattern; responses to frozen-noise patterns are reproducible over hours. NeuroDiscos thus offer a scalable route to interface and functionally characterize primary and patient-derived neuronal cultures.

## Results

### Design of the NeuroDiscos

We designed the NeuroDiscos along the four requirements outlined in the Introduction. For affordability and robustness, we used LED and OLED arrays rather than lasers and spatial light modulators; such (O)LED arrays house hundreds to millions of individually controlled emitters within a few square centimeters. In the general design, the contents of the luminous screen are controlled by a microprocessor and projected onto the light-sensitized neurons through a simple triplet lens (Fig. 1a). Various combinations of screen, recording device and upright or inverted light path are possible; here we present two. The flex NeuroDisco (flexND; Fig. 1b) projects 16 × 16 blue µLEDs through the glass bottom of a 60-electrode MEA to drive neurons expressing channelrhodopsin-2 (ChR2). The high-definition NeuroDisco (HD-ND; Fig. 1c) projects a 1,920 × 1,080-pixel green OLED display through a custom 3D-printed lid onto a CMOS MEA with 26,400 electrodes to drive neurons expressing ChReef. The lid creates an air-glass-medium interface for the projected image, while allowing gas exchange and limiting evaporation (Fig. 1d). At the culture, the devices deliver light power densities of 1-20 µW/mm^2^ (Supplementary Fig. 1).

**Fig. 1.**
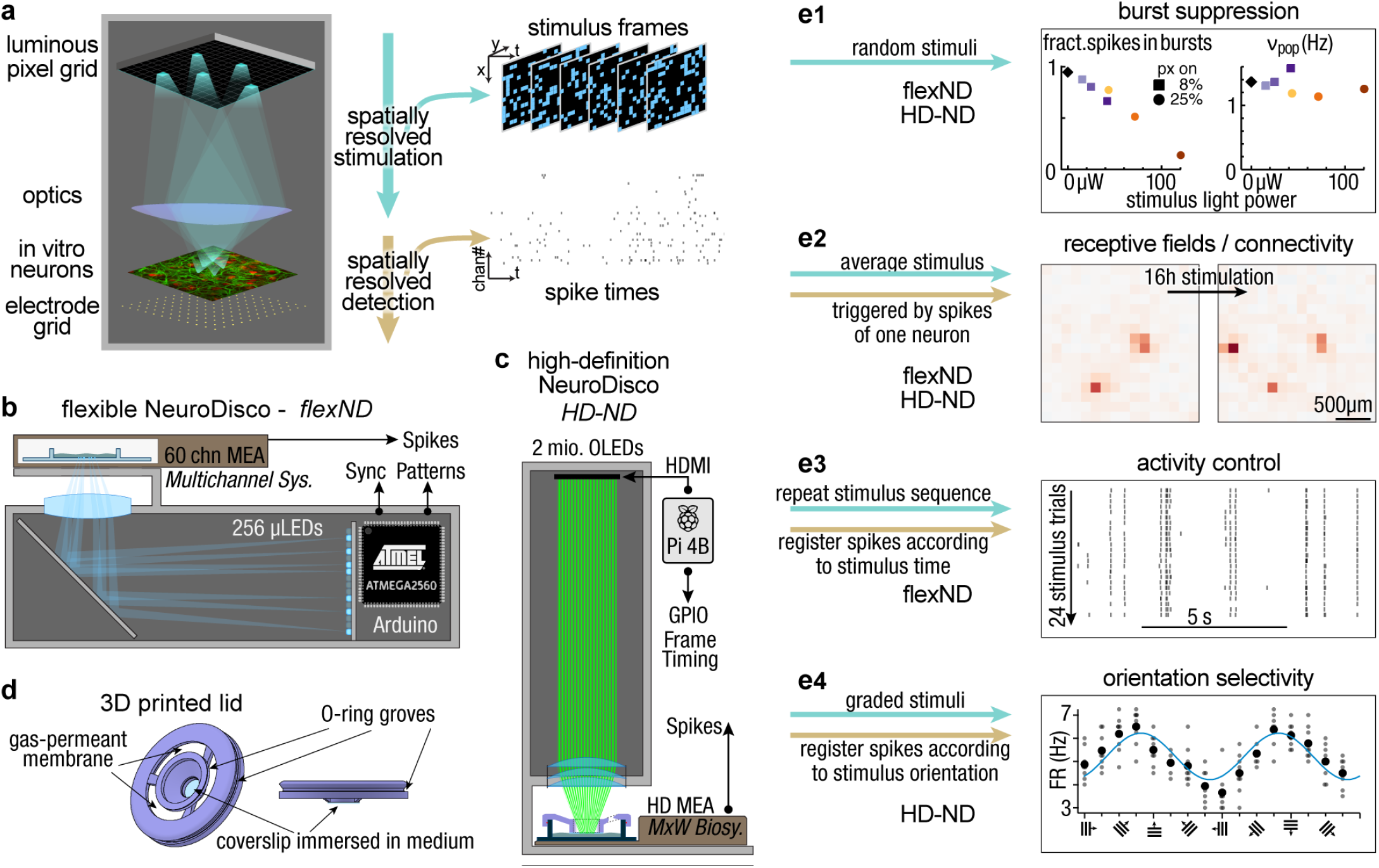
Versatile light stimulators unlock the potential of neuronal cultures. **a**, Principal design of a NeuroDisco and its output in conjunction with a synchronized recording device. **b**, Design of the flexND with an inverted light path for stimulation through a glass-bottom MEA. **c**, Design of the HD-ND for illumination through a custom-made lid. **d**, The 3D-printed lid creates an air-glass-medium interface via a coverslip. It allows for gas exchange and hinders evaporation. **e**, Utilizing patterned stimulation enables massive suppression of network bursts. **e1**, Patterned, uncorrelated stimuli disperse bursts without increasing the overall firing rate. For the experiment in Fig. 2 the average fraction of spikes that occur in bursts (left) and the average population firing rate (right) are plotted against the stimulus’ light power. The averages were taken over all time intervals that were at least 120 s after the last change of stimulus conditions. **e2-e4**, Once bursts are suppressed, various functional characterizations become feasible if stimulation is combined with synchronized electrophysiological recordings: **e2** Reverse correlation estimates receptive fields, **e3** Frozen-noise responses quantify the dynamic instability of neuronal network activity, and **e4** Responses to moving sine gratings determine the neuron’s orientation selectivity. Shown are average rates of 4 s long trials (small circles), the mean of all trials with the same stimulus direction (large circles), and a sinusoidal fit with a period fixed to 2π (blue). The labels HD-ND and flexND explain which ND type is capable of the specific experiment type.

### Synchronization with the recording

During experiments lasting days, the independent clocks of projector and recording system could drift apart by hundreds of milliseconds, too much for spike-timing analyses. We therefore devised a synchronization scheme that places stimulus events into the data-stream of the recorder and also into a data-log of the stimulator. The exact implementation depends on the capabilities of the recorder and the microprocessor. In the HD-ND, every frame update generates a digital pulse that is logged in the Raspberry Pi’s hard drive and recorded via a digital input of the MaxOne headstage (Methods). Concurrent measurements of light output and digital pulses indicate that the pulses precede the start of the screen update by 3.73 ms (median) with an interquartile range of 0.84 ms, independent of the length of the recording (Supplementary Fig. 2). In the flexND, a clock signal from the recording device serves as master for the microprocessor, which updates the screen within 7-12 µs after the trigger-signal is toggled (Supplementary Fig. 2; interquartile range 0.3 µs). The clock signal is recorded into the data-stream and its time-stamps, as registered by the microprocessor, are written to a log file, together with the content of the current frame. In both NeuroDisco systems, the redundant bookkeeping allows a complete alignment of stimulus and neuronal activity.

### Long-term operation in the incubator

For continuous operation inside the incubator, a sealed casing protects the electronics from humidity. We compensated for the heat dissipated by both the recording devices and the stimulators by lowering the incubator set point, typically to 34.5 °C, which kept the temperature at the MEA at 36-37 °C (Methods). Because blue light, unlike green light, alters activity-regulated gene expression in cultured neurons and can be phototoxic ^28,29^, we tested long-term compatibility with the blue-light flexND. Over a week of continuous stimulation, the number of spike-sorted active units did not differ between stimulated and control cultures on any of the eight sampled days in vitro (sample sizes ranged from 12 to 24 cultures per group and DIV; two-sided Mann-Whitney U tests, Holm-corrected across days; all adjusted P = 1.00; Supplementary Fig. 3).

### Spatial coverage and resolution

The flexND images its 16 × 16 µLED array onto a 2 mm × 2 mm field that covers the 1.4 mm × 1.4 mm recording area of the 60-electrode MEA. At this magnification, the image of a single µLED had a diameter of about 46 ± 2 µm (mean ± s.d., n = 256 µLEDs; Supplementary Fig. 4e); we routinely defocused it to a disc of about 90 µm, on the order of a neuron’s dendritic tree. The HD-ND illuminates the entire 3.85 mm × 2.1 mm recording area of the HD-MEA with a pixel pitch of 1.9 µm in the neuronal plane. Its resolution is limited by the lens system, which had an on-axis point-spread function of about 10 µm (from 6.9 × 9.3 µm elliptical shapes in screen corners to circular 10.8 × 10.8 µm, each given as full width at half maximum; Supplementary Fig. 4b,c). We used stimulation units of 19 µm × 19 µm, close to the 17.5 µm electrode pitch of the HD-MEA; these units stimulated neurons effectively. In both devices, all dimensions in the image plane can be scaled by moving the screen along the optical axis without compromising the light power density; the values given here were used for all experiments.

### Robustness, cost and scalability

The devices are compact (flexND: 25 × 7 × 9 cm; HD-ND: 17.5 × 16.9 × 16.5 cm) and inexpensive, with hardware costs of about €500 (flexND) and €1,000 (HD-ND), and the flexNDs have operated for years without repairs (5 devices with cumulative operating times of 5 years each). We routinely run five devices in parallel in one incubator, and up to eight can be accommodated; this limit is set by their footprint and dissipated heat. In summary, both NeuroDiscos provide spatially patterned optogenetic stimulation that is synchronized to the recording and runs inside a standard incubator.

### NeuroDiscos disperse population bursts over many hours

Theoretical work suggests that weak, irregular stimulation may be sufficient to induce asynchronous activity while maintaining firing rates in networks with homeostatic plasticity ^11^. Motivated by this prediction, we generated spatiotemporally uncorrelated patterns and applied them to cultures via the flexND (Methods). To investigate the effects of stimulation across many timescales, we recorded the activity of cultured neurons for ∼18 h, including a ∼14.3 h stimulation protocol, preceded by 1 h and followed by ∼2.5 h of spontaneous activity. During stimulation, 270 s epochs of patterned flexND illumination alternated with 270 s epochs of darkness. Each illumination epoch used one of six conditions, combining two spatial coverages (on average 8% or 25% of the LEDs switched on per frame) with three LED brightness levels, presented 16 times in a random order (Fig. 2a; Methods).

We obtained 28 of these recordings and present our findings with an in-depth analysis of a representative example (Fig. 2). The neuronal activity in this recording varied on timescales ranging from hours to milliseconds. Overall, the population mean firing rate (MFR) increased during the ∼14 h stimulation period, reaching approximately 150% of its initial baseline after stimulation had ceased (Fig. 2b,c). In addition to this long-term increase, the MFR fluctuated substantially on timescales of several minutes (Fig. 2b,c). At finer temporal resolution, synchronous network bursts dominated variability on timescales from seconds to milliseconds, with only ∼10% of spikes occurring outside these events during spontaneous activity (Fig. 2c-e).

**Fig. 2.**
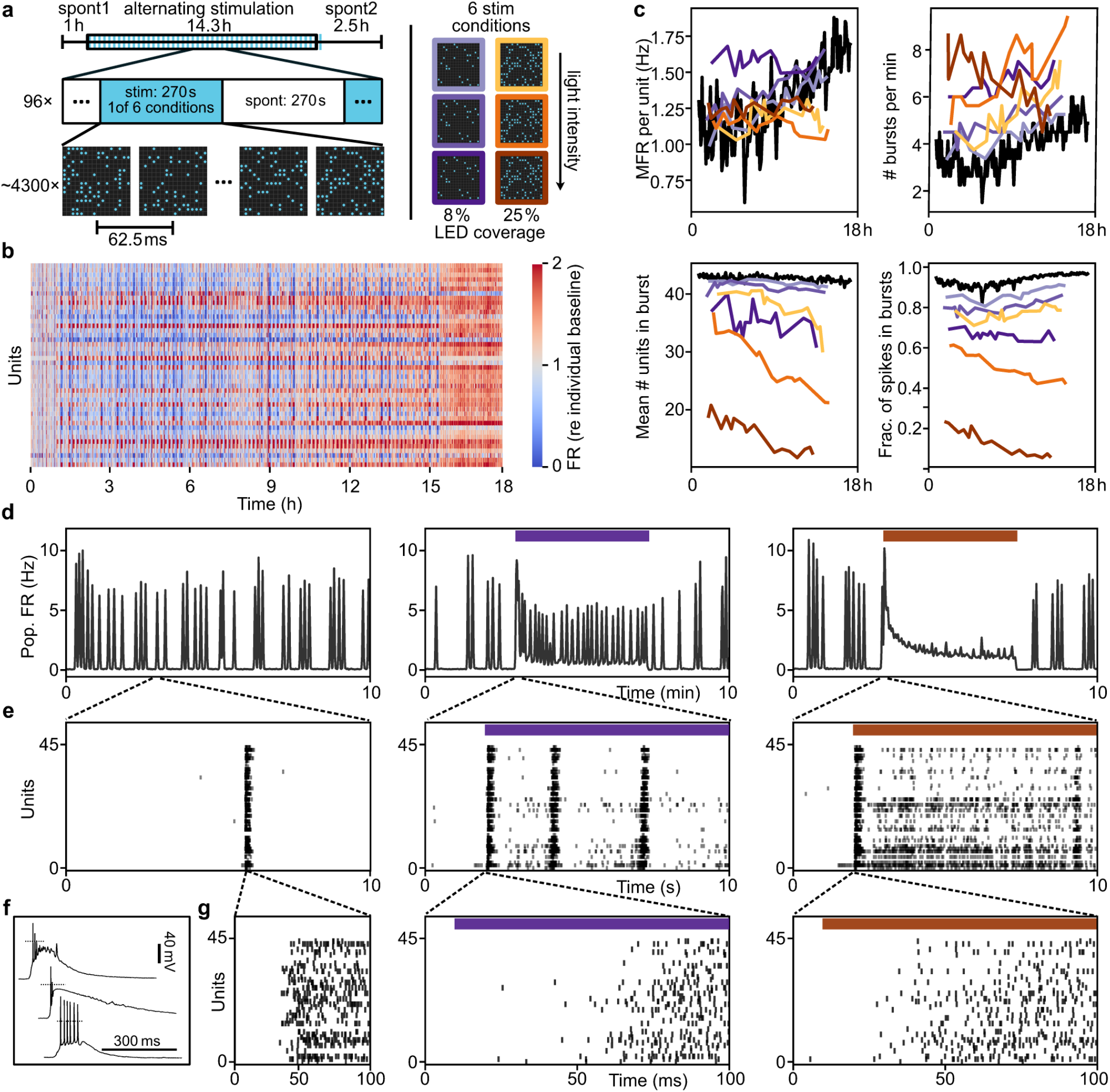
ND-stimulation modulates network properties across timescales and suppresses bursts. **a**, Scheme of experimental paradigm. Cultures were recorded for ∼18 h, ∼3.5 h without stimulation (spontaneous; sp; 1 h before and ∼2.5 h after stimulation), ∼14 h alternating darkness and light-stimulation with the flexND. Stimulation conditions varied in light intensity and spatial coverage (8% - purple shades; 25% - orange shades, details in Supplementary Fig. 1a). **b**, Heatmap of firing rate across the full 18 h recording. Each of the 45 lines corresponds to one unit (neuron). The rate is normalized to the unit’s baseline firing rate during the first hour. Bins represent 54 s. Note that the color scale is capped at 2. **c**, Network activity metrics over the 18 h recording, calculated separately for the stationary part (final 160 s) of each 270 s epoch. Data points are connected by straight lines. Colors as in a, black is spontaneous activity. **d,e,g**, Representative activity examples across three conditions (left to right: spontaneous activity; bright, 8% coverage; bright, 25% coverage). d, Population firing rate over 10 min. e and g, Raster plots of neuronal activity over 10 s and 100 ms, respectively, including the onset of a network burst. **f**, Representative traces of the membrane potential in neurons during bursts, recorded with whole-cell patch-clamp in three neurons in three cultures, different from the one shown in c-e,g. Two of the cells display paroxysmal depolarization shifts. A further 21 recordings are shown in Supplementary Fig. 7.

Besides these changes in spontaneous activity, every individual illumination epoch drove a stereotypical pattern: at stimulus onset, the MFR shot up for around a hundred milliseconds, rapidly fell and then gradually approached a quasi-stable level (Fig. 2d). Interestingly, for the population, this firing rate was close to the recording-wide spontaneous mean under both the weakest and strongest stimulation conditions, falling below it by 1.2% and 5.3%, respectively. The largest increase (+19.2%), occurred at 8% coverage and highest brightness, suggesting that the firing rate does not simply increase monotonically with overall stimulation strength (Fig. 2c; see also Fig. 1e1 and Supplementary Table 1). Individual neurons’ response to stimulation differed: some increased, and some decreased their firing rate under stimulation (Fig. 2b and Supplementary Fig. 6).

This stability in population MFR contrasted with a marked dispersion of network bursts during ongoing stimulation. Burst size, duration, and fraction of spikes in bursts decreased with stimulus coverage and illumination strength (Supplementary Fig. 5). Under the strongest stimuli, ∼90% of spikes were fired outside bursts (Fig. 2c). Bursts thus changed from the dominant activity mode in periods without stimulation to a minor modulation of ongoing irregular firing during strong stimulation.

A burst is not simply a phenomenon of particularly synchronous firing; it has drastic physiological effects on the individual neurons. Intracellular recordings revealed that bursts cause extremely strong depolarization plateaus in a majority of cells (18 out of 24 neurons from 3 preparations), sometimes reaching as high as −10 mV (Fig. 2f and Supplementary Fig. 7). Such events have been reported before in neuronal cultures ^30,31^. They resemble paroxysmal depolarization shifts observed during epileptic seizures in vivo as consequences of massive synaptic input ^8^.

### NeuroDisco stimulation shapes burst propagation away from intrinsic pathways

Population bursts are known to propagate along stereotypic trajectories ^32,33^. We captured these trajectories by ranking units according to the latency of their first spike within each burst (Fig. 3a,b; Methods). We used Kendall’s τ to quantify similarity between burst recruitment sequences (Fig. 3c) in two ways: through all-to-all pairwise comparisons and by comparing individual bursts with a reference pattern. The reference pattern was defined by the median recruitment rank of each unit across bursts during spontaneous activity recorded approximately in the first 5 min (Methods). Across the 18 h recording, most spontaneous bursts followed similar recruitment sequences, with 83.6% exhibiting Kendall’s τ ≥ 0.5 relative to the reference pattern (Fig. 3d,e,g). However, distinct subclusters of bursts were also present, including bursts that showed negative correlations with the reference pattern (Kendall’s τ < 0; 7.0% of spontaneous bursts, Fig. 3d,e,g). This might reflect a reversal of the typical initiation sequence.

Given that stimulation dispersed bursts, we examined whether the remaining bursts during stimulation still shared those intrinsic routes or initiated along altered trajectories. We compared their trajectories again with the spontaneous reference pattern (Methods). Across stimulation strengths, most bursts showed positive correlations with the reference pattern (Fig. 3e). Even in the condition showing the strongest effect (25% coverage and highest brightness), this proportion remained at 82.2%. However, in this condition, the proportion of strongly correlated bursts (Kendall’s τ ≥ 0.5) decreased from 83.6% during spontaneous activity to 29.1%, while the proportion of negatively correlated bursts (τ < 0) increased from 7.0% to 17.8% (Fig. 3e,h). Stimulus-onset-induced bursts showed no correlation with the reference pattern (Fig. 3f).

**Fig. 3.**
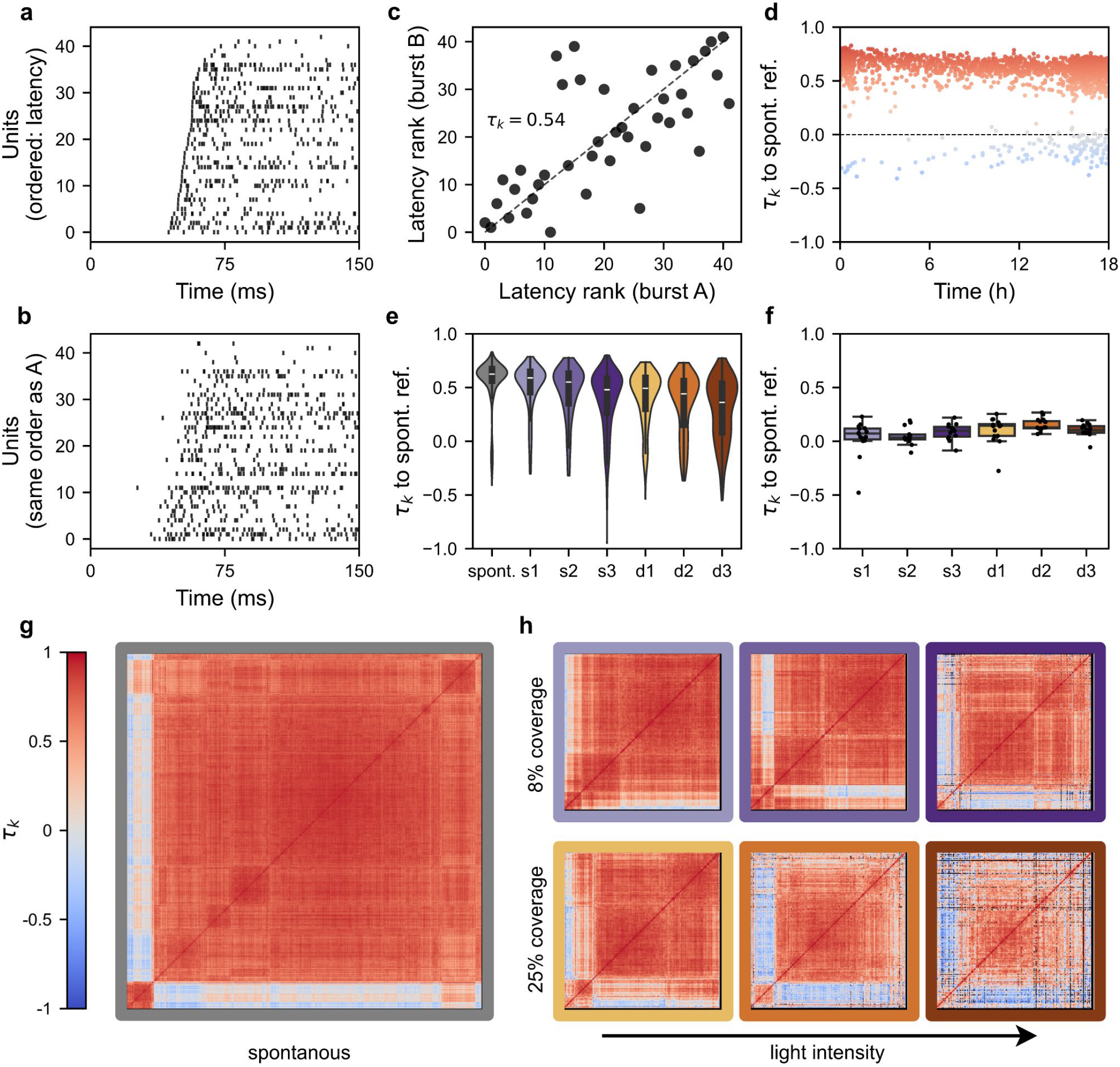
ND-stimulation acutely destabilizes intrinsic propagation trajectories. **a**, Raster plot of a burst during spontaneous activity approximately 10 min into the recording. Units are ordered by their first spike latency during this burst. **b, As in a**, but for a burst recorded 17 h later, same unit order. Note the similar initiation sequence. **c**, Scatter plot of the ranks of all units in the two bursts from a and b. The dashed line indicates identity. The strong alignment between the ranks in bursts 17 h apart is represented by a high Kendall’s tau (τ_K_). **d**, Kendall’s tau plotted for each burst in the spontaneous periods, excluding the first 110 s after each stimulation period. A reference ranks vector was constructed from the median rank for all bursts during the first 276 s (Methods). Note the small set of bursts with negative correlation (blue) (7% of spontaneous bursts), indicating propagation along the reference route but in reversed order. **e**, Violin plots showing Kendall’s τ between individual burst trajectories and the spontaneous reference pattern during light-driven (colored) and spontaneous (gray) activity. Stimulation conditions are ordered by increasing stimulus strength and coverage (8%: s1-s3; 25%: d1-d3). Burst counts were n = 194, 209, and 268 for s1-s3, respectively; n = 233, 273, and 230 for d1-d3, respectively; and n = 1,902 for spontaneous activity. Colors as in h and Fig. 2. **f**, Tukey boxplots showing Kendall’s tau as in **e**, but restricted to the first burst of each illumination period (n = 16 bursts per condition). **g**, Matrix of Kendall’s tau values for pairwise comparisons of 1,902 spontaneous bursts. The order results from similarity clustering, and reveals the anti-correlated subgroup. **h**, Matrices of Kendall’s tau values as in g, but for the bursts during light-driven activity (bursts from **e**) and with black assigned to comparisons that would be based on fewer than five shared units (Methods). All data are from the recording in Fig. 2.

### NeuroDisco-derived receptive fields monitor neuronal encoding and functional connectivity

Spike-triggered averaging originated in the white-noise analysis of sensory systems ^34^ and has become a standard, conceptually simple tool for characterizing neuronal encoding ^35–37^. The NeuroDisco’s precise registration of every stimulus frame to the recording allowed us to apply it to cultured networks: we computed spatiotemporal receptive fields (RFs) as the spike-triggered average (STA) of the stimulus preceding each recorded spike. Patterned holographic stimulation has been used in this way to map the stimulation fields of MEA-recorded units ^25^; here, mapping covers the entire recording area and continues for many hours.

Already a few hundred spikes, typically acquired within 3 min of driven activity, were sufficient to localize the RF (Supplementary Fig. 8a). With increasing numbers of spikes finer details become discernible (Supplementary Fig. 8a). A typical RF displayed a dominant spatial component extending over a few hundred micrometers around the recorded unit, a scale compatible with local somatodendritic contributions ^38^. In some cases, the high resolution of HD-ND even revealed elongated RF components that might represent dendritic structures (Fig. 4b,c). About a third of the receptive fields showed biphasic temporal profiles (32 of 100 units, from four recordings), with an early negative phase followed by a positive phase closer to spike time (Fig. 4a). Similar biphasic stimulus-response kernels were observed in patch-clamp experiments ^39^ and likely reflect an increase in neuronal excitability after sodium channels de-inactivated at less depolarized membrane potentials.

**Fig. 4.**
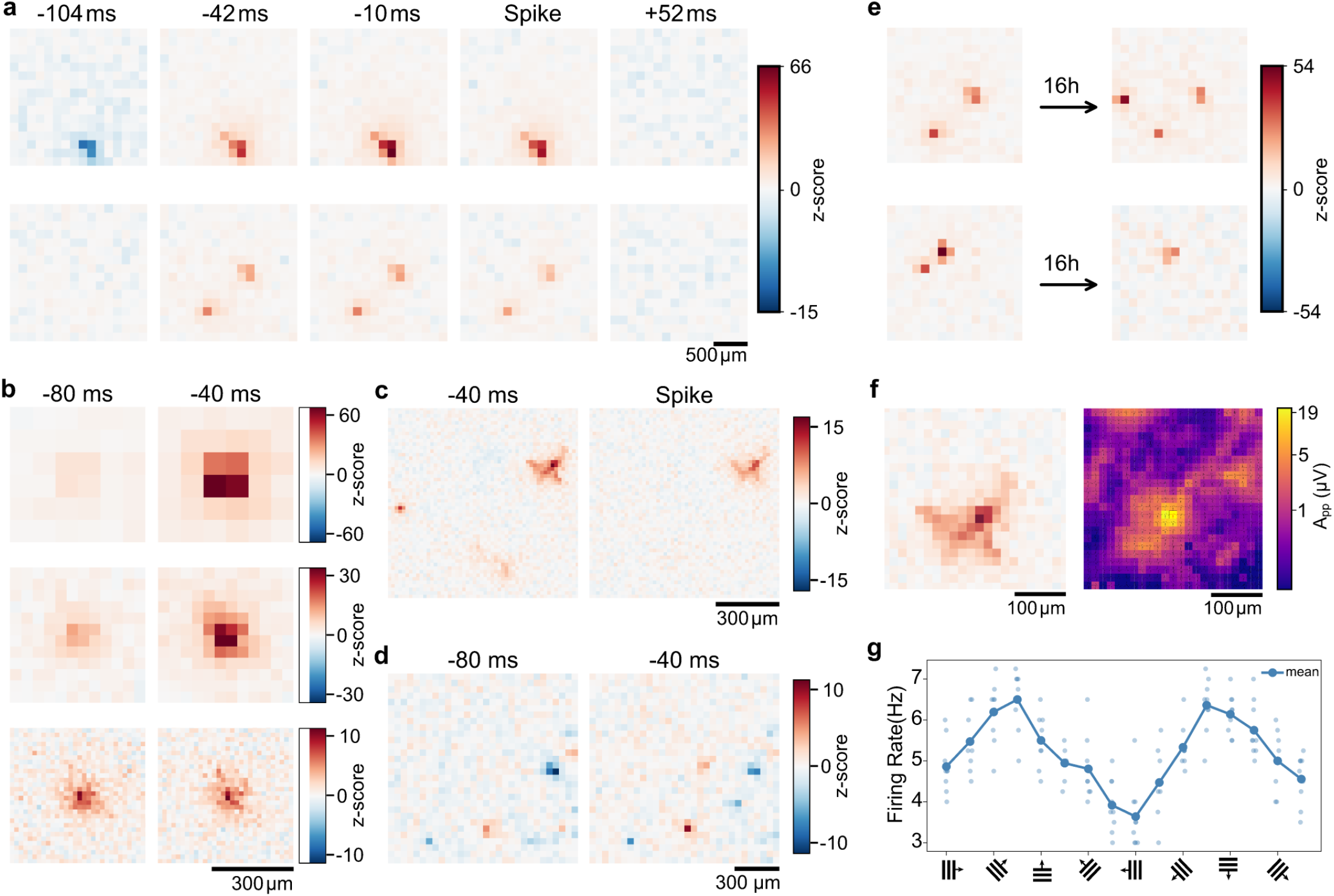
Receptive fields reveal stability and plasticity of cellular components and network interactions. **a**, Two representative spatiotemporal receptive fields (RF), calculated as spike-triggered average stimulus patterns, for two units recorded during Poisson stimulation with the flexND (Methods). Note the blue shadow at 100 ms before the spike. **b**, Receptive fields of one neuron obtained using HD-ND Poisson stimulation patterns with 92, 47.5, and 19 µm pixel sizes (15 min each). Note that the lowest resolution is comparable to the 136 µm resolution of the flexND. **c,d**, Examples of complex negative and positive subfields in RFs. Note the different temporal dynamics of the distinct subfields. **e**, RFs of two cells obtained under Poisson stimulation before and after 16 h of correlated stimulation (Methods) show substantial changes in RF structure. **f**, Comparison of the RF and the spike-triggered extracellular potential around the same neuron. Color code encodes the logarithm of the peak-to-peak difference of the extracellular potential in a 4 ms bin, centered on the local minimum of the waveform of the triggering electrode (red waveform). **g**, Orientation tuning curve obtained from 9 runs of 16 orientations of sine gratings with 760 µm wavelength. Supplementary Fig. 8 documents RF stability and spike-count dependence.

To further investigate the nature of the RFs, we produced a spike-triggered electrical signal from electrodes surrounding the triggering unit and compared it with this unit’s RF (Fig. 4f). The apparent similarities between these two maps strongly support the hypothesis that the central component of the RF reflects dendritic structures. The shape of the extracellular action potential waveforms in this area also supports the dendritic nature, in particular their strong initial depolarization and the peak latency of about 1 ms.

The RFs of some neurons contained additional components outside the plausible dendritic field, up to 1 mm away from the recorded unit (Fig. 4a,c-e). In the recordings analyzed here, 20% of the units with a receptive field carried such a separate component (21 of 106 units from four recordings). These subfields can be positive (Fig. 4a,c) or negative (Fig. 4d) and likely capture activation of excitatory or inhibitory neurons, which affect the triggering neuron. We conclude that, at least for particularly strong interactions, NeuroDisco-derived RFs are able to detect functional connections between neurons or local clusters.The RFs of many neurons remained stable over hours. In a continuous recording with uncorrelated stimulation alone, the same RF could be recovered throughout the recording (Supplementary Fig. 8b). We also assessed RF stability in four recordings in which 16 h of spatially correlated stimulation (Methods) were preceded and followed by 1 h of fully uncorrelated Poisson stimulation. RFs were estimated separately from these two Poisson stimulation periods. Of the 99 units with a detectable RF during the first hour, 72 (73%) still had a detectable RF after correlated stimulation, whereas RFs were no longer detectable in the remaining 27 (27%). Among the 72 units with RFs detectable at both time points, the location of the dominant pixel was unchanged in 57 (79%). A small subset of units showed distal RF components that appeared or disappeared between the two measurement periods (Fig. 4e). These changes in putative connectivity showed no obvious relationship to the spatial correlation structure of the stimuli.

The receptive field of a neuron can not only be characterized in the spatial domain, but also in the Fourier representation, the k-space. Stimulation with sinusoidal waves of a particular spatial frequency belongs to this kind of characterization. Utilizing the HD-ND’s grayscale screen, we presented moving gratings of 16 different directions to the neurons. A small fraction of cells showed very clear orientation selectivity (Fig. 4g), presumably because their RF had a strongly asymmetric shape.

### flexND tames the dynamic instability of neuronal cell cultures

Repeated applications of the same stochastic stimulus (frozen noise) test how reliably a system encodes various input features. Individual cortical neurons fire very reproducibly in response to fluctuating input ^40^. Inside networks, the input-output mapping may be less reliable due to chaotic dynamics, internal noise, and slow drifts of the network state^41^. Such experiments require many repetitions under stationary conditions and many spikes per neuron. Because NeuroDiscos allow for precise stimulation over many hours inside the incubator, we could present frozen-noise sequences up to 96 times (4 intensity levels × 24 repetitions) over ∼15 h and record 10^5^-10^6^ spikes from individual units. Here we use these data to show how reproducibility depends on stimulus strength and network state; a quantitative analysis of the underlying dynamics is beyond the scope of this paper.

The overall experiment design corresponds to Fig. 2a, with some changes to the duration and number of the stimulus and dark periods. Importantly, stimulation patterns are created from random processes with known random seeds. This allowed us to produce exact repetitions of two pattern sequences, presented at four different light power levels, determined by the LED controller’s voltage (Fig. 5a and Supplementary Fig. 1a). The spike sequences that a given unit produces under repetition of the same stimulus were remarkably similar, even if the trials ran 10 h apart (Fig. 5b). Similarity, as measured by the spike time tiling coefficient (STTC) ^42^, increased with light power (Fig. 5b-d). Unlike the sequence similarity in spontaneous bursts (Fig. 3), the similarity under stimulation does not result from imprinted propagation pathways. Instead, the spike sequence follows the stimulus pattern, i.e., a different stimulus pattern sequence leads to completely unrelated spike sequences (Fig. 5d). Whenever network bursts occur, spiking is dissociated from the external input and reproducibility collapses for several seconds (red marks in Fig. 5e).

**Fig. 5.**
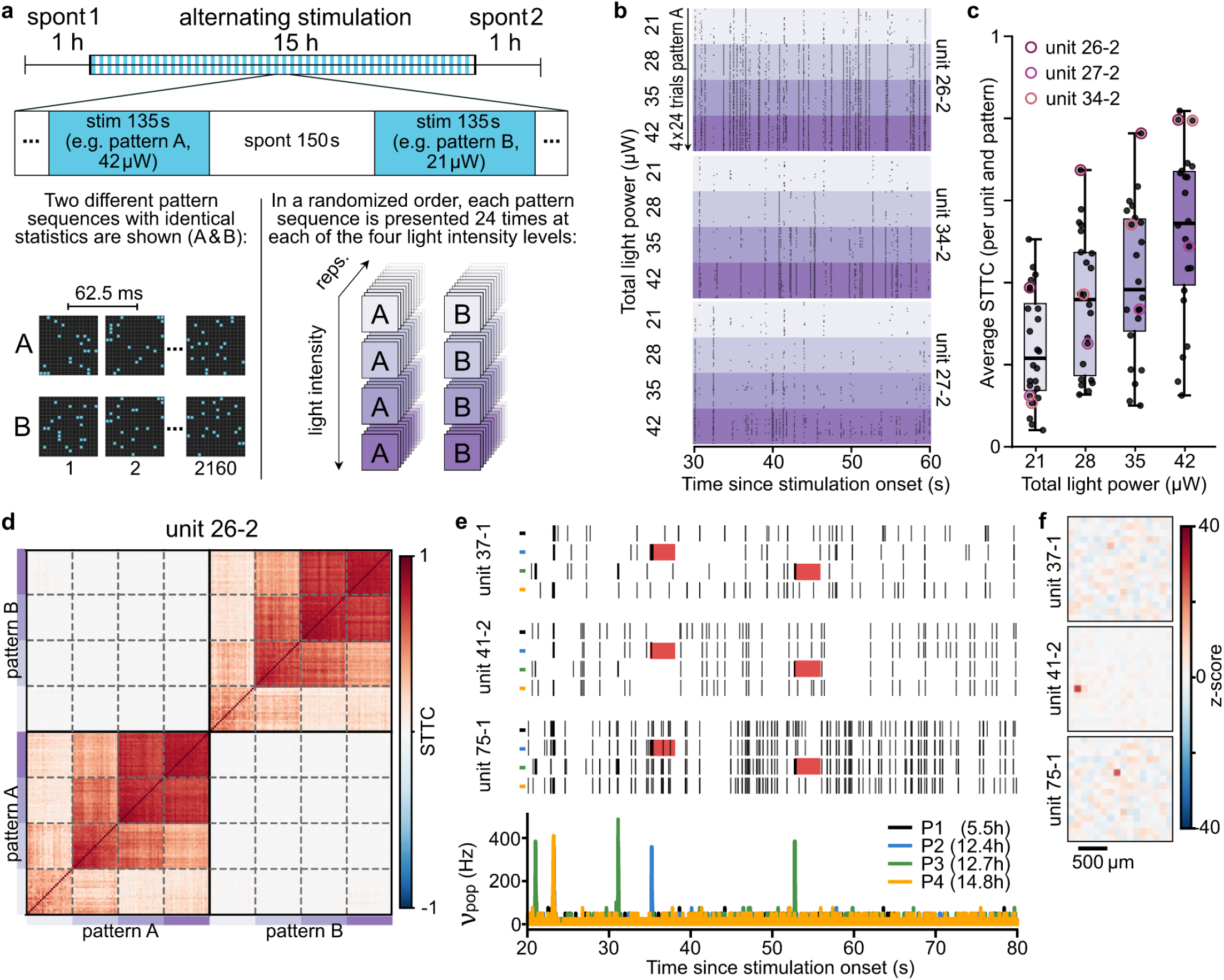
Weak stimulation established control over dynamic population activity. **a**, Scheme of the experimental paradigm. For details, see Methods. **b**, The spike raster plots for three units in response to identical stimulus sequences (pattern A). In each colored block, the responses to 24 trials with identical light powers appear in chronological order from top to bottom, spanning the 15 h stimulation block (for light power details, see Supplementary Fig. 1a). **c**, STTC is used to quantify the similarity between spike sequences fired in response to identical patterns presented at identical light power. Results from the data in b are highlighted by circles in red-purple shades. **d**, All-to-all STTC similarities for all 192 responses of unit 26-2 for all four light powers and both pattern sequences. Within each block, responses are arranged in temporal order. a-d show the design and results from one representative recording out of 20 we conducted with identical design. **e**, In an earlier experimental design, 4 repetitions of one stimulus pattern were presented for each of three different light powers. Shown are the responses of three units to pattern A presented with 42 µW light power. The bottom panel shows the MFR during these four stimulus repetitions. Note the large MFR transients that correspond to bursts. The burst at 23 s is precisely reproduced in every trial. Other bursts occur only in a single trial and transiently reduce the response precision, indicated by the red marks. **f**, The RFs for the three units in e. Note that unit 37-1 appears not to be modulated by light stimuli.

While the increased response reproducibility under stronger external drive is expected, it is intriguing that this dependence is observed in all units (Fig. 5c). We can be certain that some cells do not express ChR2 and are thus not light-sensitive. To investigate this further, we calculated the RFs of all units. We found that units with strong RFs typically produce sequences with narrow, strong cross-correlations. But even units with no discernible RF and thus with negligible light-sensitivity produced correlated spike trains on repeated trials (Fig. 5e,f). This suggests that direct stimulation of light-sensitive units also reliably recruits other units into the network activity, even if these are not light-sensitive themselves. We are thus capable of controlling and quantifying the regularity of the collective activity even beyond the directly illuminated area.

## Discussion

The NeuroDiscos are compact, affordable projectors that deliver long-term, spatially structured optogenetic stimulation to light-sensitized neurons inside a standard incubator. They are synchronized to concurrent extracellular recordings with submillisecond precision that is maintained over recordings lasting days to weeks. We used this capability for two purposes: to place cultures in a sustained asynchronous activity regime, and, in that regime, to characterize the encoding and connectivity of individual neurons.

Patterned optogenetic stimulation with µLED and OLED arrays offers several advantages over existing ways of stimulating neuronal cultures. Unlike electrical stimulation, optical stimulation does not interfere with the ongoing recording and offers greater flexibility with its hundreds to thousands of spatially addressable, concurrently operating optical stimulation elements. Optogenetic stimulation of cultures has so far followed two routes. Whole-field LED illumination ^20–22^ provides no spatial structure. Microscope-based pattern illuminators – confocal scanners, digital micromirror devices (DMDs) and holographic systems – are costly and typically operate on the stage of a large microscope rather than in the incubator; reported sessions have lasted only minutes to about an hour at irradiances of 5-15 mW/mm^2 43,44^. The NeuroDiscos combine the strengths of both routes: like whole-field LED systems, they are inexpensive, scalable, and run continuously in a humidified incubator for days to weeks; like microscope-based systems, they deliver spatially resolved patterns. The 19 µm stimulation units of the HD-ND are finer than the 50 × 50 µm spots recently used for single-neuron stimulation with a DMD on the same HD-MEA platform ^44^, and the ∼90 µm spots of the flexND are comparable to those of an earlier DMD-based culture stimulator (∼100 µm) ^43,45^.

Remaining technical limitations concern the temporal and spatial resolutions. The frame rate is ∼200 Hz for the flexND and 22.5-60 Hz for the HD-ND. Spike timing precision is further limited by the low light intensity that leads to longer latency and larger jitter. Thus, the current NeuroDisco versions are adequate for graded modulation of spike timing but too slow to precisely modulate activity on the timescale of monosynaptic delays. The spatial resolution is bounded by the ∼10 µm point-spread function of the budget optics. In the flexND, the relatively large emitter size and pitch also limited the resolution. These issues will be addressed by software and hardware updates. Specifically, Cooke triplet lenses should reduce optical aberrations, and dedicated LED controllers can dispense with row multiplexing and thereby speed up pattern delivery and increase the light power density at least fivefold. In addition, future ND hardware could combine two screens of different colors via a dichroic mirror and drive spectrally separated opsins, for example, to add an inhibitory channel.

All optogenetic stimulation carries a risk of phototoxic effects, particularly with long-term use. Even at the low irradiances produced by the NeuroDiscos, the blue light of the flexND has the potential to induce phototoxic products in riboflavin-containing media such as Neurobasal/B27 ^28,29^. Although our long-term tests showed no indication of cell loss (Supplementary Fig. 3), photo-inert media or green-light stimulation (HD-ND) should be preferred in transcription- or plasticity-sensitive experiments.

The NDs offer one key advantage for studies of neuronal physiology and information encoding: they can disperse the spontaneously occurring population-wide bursts. These bursts comprise as much as 90% of all spikes, drive more than half of the cells into depolarization block (18 out of 24; Fig. 2f, Supplementary Fig. 7), and behave as near all-or-none events ^7^. While a burst unfolds, spiking is decoupled from the external input, and stimulus-locked reproducibility collapses for several seconds (Fig. 5e), and responsiveness recovers only gradually afterwards ^6,46–48^. Such strong, lasting depolarizations likely interfere with normal synaptic signaling and plasticity. Stimulating with spatiotemporally decorrelated patterns substantially dispersed the bursts over hours and days, but left the firing rate essentially unchanged. This dissociation of decorrelation from firing rate distinguishes our approach from drive-based burst suppression, which raises tonic firing and has been sustained only for minutes ^49^. Because the random pattern stimulation keeps the population in an asynchronous regime where spike timing is shaped by the external input (Fig. 5b-d) and by the ongoing network state ^50^, we posit that NDs could improve the physiological validity of a wide range of studies in cultures.

To illustrate the range of possible applications of the irregular activity, we exemplified functional characterizations of individual cells and the population’s dynamic regime. Spatiotemporal RFs can be determined within a few minutes (Supplementary Fig. 8a). They indicate the cells’ location in stimulus space, their light sensitivity, and, in some cells, putative connections to excitatory or inhibitory partners up to a millimeter away. This information can be used to plan cell- and connection-targeted stimuli in the same culture just minutes later. At the HD-ND’s resolution, RFs also revealed elongated and multi-component structures consistent with somatic and dendritic contributions. The networks’ dynamic stability was characterized with repeated frozen-noise sequences, which showed that stimulus-locked spike trains are reproducible across trials ten hours apart, and that reproducibility increases with light power. Because all recordings ran in a humid incubator for many hours under quasi-stationary conditions, we could also observe slow drifts of firing rate, burst propensity, and RF structure.

The NeuroDiscos open new paths for basic and translational research in vitro. Pharmacological screenings and phenotype characterization can be performed in a more physiological asynchronous regime, without exposing the neurons to pathological depolarizations. The most immediate application is the study of plasticity. Because cultures can be held in an asynchronous regime for days while arbitrary spatiotemporal stimuli are imposed and responses are recorded continuously, any conditions that induce, maintain or reverse network changes can be varied systematically in the same culture. Signatures of such changes are already apparent in our recordings: the population firing rate reached ∼150% of its initial baseline after ∼14 h of patterned stimulation, and in a small subset of neurons distal receptive-field components appeared or disappeared over 16 h of spatially correlated stimulation. Delivering correlated stimuli without the unphysiologically strong depolarizations of bursts may permit a more physiological, spike-timing-dependent regulation of synaptic strength and longer-lasting synaptic memory than is typical in vitro. Beside the characterizations presented here, the asynchronous regime should make correlation- and information-based measures more informative, which in turn deepen analyses of functional connectivity and information encoding. Closing the loop would then allow stimulation protocols to be optimized rather than fixed in advance: a coarse receptive field is available within minutes so stimulation can be confined to a neuron’s identified presynaptic territory or depend on prior spiking ^51^.

The same capabilities carry over directly to patient-derived neurons. Functional assays of such cultures currently rely largely on population-burst statistics, which may mask subtle but functionally meaningful alterations. Holding patient-derived networks in an asynchronous regime and probing them with defined stimuli would allow genetic pathologies to be characterized at the level of single-neuron encoding, connectivity and network stability, and their rescue by pharmacological treatment or genetic correction to be followed in the same culture over days. Drug effects could likewise be assessed on stimulus-driven responses rather than on spontaneous bursting alone; repeated frozen-noise stimulation, for example, provides a direct readout of how a compound or mutation alters the dynamical stability of the network. At hardware costs of approximately €500 (flexND) and €1,000 (HD-ND), the provided construction files make this technology widely available and allow experimental scale-up, especially in combination with emerging open culture-and-recording hardware ^17^. Further miniaturization of the projectors to multi-well MEA formats would provide the throughput required for medium-throughput screens. We anticipate that bringing spatially resolved, incubator-stable, long-term optical control to standard culture platforms will make functional phenotyping a routine complement to the molecular and genetic characterization of neuronal cultures.

## Methods

### Projectors

#### Lenses

Both NeuroDiscos projected their screen onto the culture with the same three-lens system: two plano-convex lenses (f = 40 mm; Linos G31 2314 000, Excelitas Technologies) and in between one concave-convex lens (f = 50 mm; Linos G31 5505 000). The lenses are mounted with a free aperture of 21.4 mm.

#### Optical pathway

In the flexND the distance between LED screen and lens (planar end) is 23 cm, the distance between lens and image approximately 1.7 cm. This system demagnifies by a factor of 0.071. An aluminum-coated glass mirror (40 × 30 × 1.3 mm; 506.VFG, Astromedia) was positioned 20 cm from the lens at 45° to the optical axis. The mirror redirected the light path so that the culture was illuminated from below through the glass bottom of the MEA. In the HD-ND the distance between OLED screen and lens (planar end) is 64 mm, the distance between lens and image approximately 16 mm. This system demagnifies by a factor of 0.34.

#### flexND

The screen of the flexND was a 16 × 16 array of blue surface-mount LEDs (19-213/B7C-AQ2S1B2/3T, Everlight Electronics; emission peak 470 nm; see Supplementary Material), controlled by a microcontroller (ATmega2560-16AU, Microchip Technology). The microcontroller drove the LED matrix using two eight-channel UDN2981 source drivers (Allegro MicroSystems) to supply current to the selected LEDs and two eight-channel ULN2803 Darlington sink drivers to provide the return path to ground. Global brightness was set by the supply voltage of the LED array (Supplementary Fig. 1), provided by a laboratory power supply (PL303QMD-P, Aim-TTi). Electronics and optics were enclosed in a sealed aluminum casing of 250 × 68 × 90 mm (L × W × H; 110 mm wide with optional cooling fins). The flexND is paired with the MEA2100-Mini-System (Multi Channel Systems) and a data logging PC. The central hub is a relay box that connects to the PC via USB, to the MEA system with 50 Ω coaxial cables (RG174 A/U) with LEMO 00 plugs (FFA.00.250.NTAC31, LEMO), to the laboratory power supply via banana plugs (LED control voltage), and to the flexND via a D-Sub cable and plug. The relay box is powered by a conventional 5 V power supply. The custom D-sub cable combines the USB connection (PC-microprocessor), the 5 V microcontroller power, the LED controller supply voltage (0-6 V) and one TTL bus (MEA system digital output to microprocessor pin PCINT1) (Supplementary Material: Controller.pdf). To store the outgoing TTL pulses in the MEA system’s data stream, they are looped back to the digital in of the recording system using the same kind of LEMO cable split using a LEMO Y-splitter (FTY.00.250.NTF).

#### HD-ND

The screen of the HD-ND was a 0.49-inch green OLED microdisplay (SY049LDM02, SeeYA Technology; 1,920 × 1,080 pixels of 5.616 µm × 5.616 µm, active area 10.783 mm × 6.065 mm, emission peak 539 nm, nominal maximal luminance 20,000 cd/m^2^, 60 Hz refresh rate). Its MIPI interface was connected to an HDMI driver board (5B0049H04-1, Shanghai Top Display Optoelectronics), which was driven by a single-board computer (Raspberry Pi 4 Model B, Raspberry Pi Ltd) operated headless via SSH over Ethernet. Brightness and power are provided via the HDMI and USB connection from the Raspberry Pi. Electronics and optics were supported by a polylactic acid (PLA) casing with outer dimensions of 175 × 169 × 164 mm (L × W × H). The MIPI interface and HDMI driver board are contained in a polystyrene Petri dish glued to the casing. The image was projected onto the HD-MEA from above through a coverslip (10 mm, No. 2, borosilicate glass) in a custom lid. The coverslip formed an air-glass-medium interface (Fig. 1d), and a gas-permeable membrane allowed gas exchange (ALA MEA-MEM-SHEET). The lid was 3D-printed from PLA using an H2D AMS Pro printer (Bambu Lab) and placed directly on the MaxOne culture well. It contained two silicone (MVQ) O-rings (38 × 1.5 mm, part no. 401070-3-21646-6400131; 18 × 1 mm, part no. 401070-5-104816-3500101, Fig. 1d). Before use, the assembled lid was treated with ethanol for 30 min and rinsed three times with sterile Type I water.

#### Stimulus creation, presentation and synchronization with the recording system

The master clock was set by the recording system; it toggled a TTL signal for every frame switch. Each pulse invoked an interrupt service routine that replaced the active pattern with the pending one, time-stamped the switch and sent the new pattern with its timestamp to a PC via a serial connection using a custom Python script. A separate interrupt service routine, invoked every 288 µs, reads the active pattern and controls the illumination of the LEDs, multiplexed line by line. Between the interrupts the main routine computed the next pattern and stored it as pending.

The new frame reached half-maximal intensity within 7-12 µs of the TTL toggle (median delay: 6.97 µs; interquartile range: 0.26 µs; Supplementary Fig. 2a). Offline, each logged pattern was matched to its TTL event in the time base of the recording. Because clock signals were logged in both devices, the data stream and the microprocessor’s log, we could catch rare cases of missed pulses (about 1 in 58,000, i.e. 1 per hour) and correct for them.

For the HD-ND, custom C code on the Raspberry Pi wrote each new frame into the screen buffer and issued a present call to the GPU, which returned after the onset of the next vertical synchronization (vsync). The code then emitted a 200 µs, 3.3 V pulse on a general-purpose input/output (GPIO) pin (BCM 27, pin 13). The pulse reached digital input D0 of the MaxOne recording unit (MaxWell Biosystems) through a Dupont jumper cable joined to the 50 Ω coaxial cables (RG174 A/U) with LEMO 00 plug (FFA.00.250.NTAC31, LEMO). Each frame switch was thus recorded twice: in the MaxOne data stream, and together with the frame number in a log on the Raspberry Pi. This allowed stimulus and recording to be aligned offline. Timestamp jitter was a few microseconds on both devices; the screen update began 3.73 ms after the vsync signal (median; interquartile range 0.84 ms; Supplementary Fig. 2).

#### Light Power and point spread function measurements

The light power was determined with a calibrated photodiode (Hamamatsu S2386-8K) in series with a low-resistance current meter. In that configuration the producer-provided calibration sheet states a conversion of μA/mW at 473 nm and 353 μA/mW at 539 nm. The total light power was measured while a fixed Bernoulli-pattern was focused onto the photodiode’s active surface. For these measurements, we used a random pattern, in which 20, 64 or 256 of the 256 LEDs of the flexND were active and determined the photocurrent for various LED controller voltages in steps of 0.1V. For the HD-ND, we chose a pattern with 10% and 100% of blocks at maximal brightness. The light power values obtained in that way are shown in the supplementary figure 1a. Upon re-measurement with a PM100D photometer with S120C sensor (Thorlabs), those values were confirmed within 5% deviation.

The images of the stimulation patterns were also projected directly onto the Sony IMX477R sensor of a Raspberry Pi HD camera. This chip has a pixel pitch of 1.55µm x 1.55µm. These images (supplementary fig. 4) were used to determine the spatial extent of the point spread function and for the estimation of the local density of the light power in the images of the LEDs.

### Cell Culture

All procedures were performed in accordance with German law on animal research and registered with the animal protection offices of the University Medical Center Göttingen and the University of Göttingen under the licenses T 23.13 and T2-2025. We modified the cell culture preparation protocol from ^52^. Newborn mice (postnatal day 0) were sacrificed by decapitation, and their brains were extracted and processed in cooled Neurobasal medium with HEPES buffer. The cortices were isolated, sparing the subcortical areas, and processed in batches of littermates. To dissociate the cells, the cortices were incubated in 0.05% Trypsin/EDTA (Thermo Fisher, #25300054) at 37 °C for 10 min, followed by mechanical trituration in Neurobasal medium with 1% FBS (Sigma-Aldrich, #F7524). After centrifugation, the cells were resuspended in Neurobasal medium containing Glutamax (Thermo Fisher, #11574466) and B27 (Thermo Fisher, #15717988), and filtered through a 70 µm cell strainer (Thermo Fisher, #11597522). Cell density and viability were assessed using an improved Neubauer hemocytometer after Trypan blue staining. To promote homogeneous transduction, the viral vector was added directly to the cell suspension before seeding at a nominal dose of approximately 1.2-4 × 10^4^ viral particles per counted neuron. The cell suspension, at a density of either 3,000 or 8,000 cells/µl, was seeded onto the electrode area in 50 µl aliquots. After 2 h, the MEAs were filled with Neurobasal medium, supplemented with Glutamax and B27.

Medium exchange was performed twice weekly. To minimize evaporation effects, MEAs were weighed on the day of cell culture preparation and at each medium exchange. The weight difference indicated the amount of evaporated water and was replenished during each medium exchange with sterile Type I water.

#### Coating

The following procedure is adopted from the preparation guide by MaxWell Biosystems. Primary coating of the microelectrode arrays (MEAs) was performed using 0.07% polyethylenimine (PEI; Sigma-Aldrich, P3143) diluted in 1X borate buffer. A volume of 1 ml of PEI solution was applied to 60-electrode MEAs, while CMOS MEAs received 50 µl. Both types of MEAs were incubated at 4 °C for a minimum of 12 h. Following incubation, the PEI solution was aspirated, and the dishes were washed four times with sterile Type I water. After drying, the electrodes were covered with 50 µl of laminin (0.02 mg/ml; Sigma-Aldrich, L2020) and left to incubate at room temperature for at least 30 min. The laminin solution was aspirated immediately before cell seeding, without subsequent washing.

#### Expression of channelrhodopsin variants by viral transduction

To enable optogenetic stimulation, neuronal cultures were transduced with one of two AAV2/9 vectors encoding distinct channelrhodopsins. Both vectors comprised an AAV2 vector genome packaged in an AAV9 capsid and used the human synapsin promoter to drive transgene expression predominantly in neurons.

The ChR2 viral preparation was supplied by the Viral Core Facility of Charité - Universitätsmedizin Berlin (RRID). The transfer plasmid, pAAV-hSyn-hChR2(H134R)-EYFP (deposited by Karl Deisseroth; Addgene plasmid #26973; RRID), encoded the H134R variant of humanized channelrhodopsin-2 fused to enhanced yellow fluorescent protein (eYFP).

For ChReef expression, cultures were transduced with AAV2/9-hSyn-ChReef-TS-eYFP-ES-WPRE-SV40 poly(A), which encoded ChReef fused to eYFP and included trafficking and endoplasmic reticulum export sequences ^53^. The vector also contained the woodchuck hepatitis virus post-transcriptional regulatory element (WPRE) and an SV40 polyadenylation signal.

### Multi-electrode Recordings

All recordings were conducted inside an incubator with 5% CO_2_ and 95% humidity. Electrical activity on 60-electrode MEAs was recorded with the MEA2100-Mini-System (Multi Channel Systems, Reutlingen, Germany). We either used 60MEA200/30iR-ITO or 60PedotMEA200/30iR-Ti (200 µm electrode pitch, 30 µm electrode diameter). Data were obtained with the Multi-Channel Experimenter Software (Multi Channel Systems, Reutlingen, Germany), a sampling rate of 50 kHz and was filtered with a hardware high-pass filter at 1 Hz and a low-pass filter at 3.5 kHz. To maintain physiological temperatures (36-37 °C) during and between recordings, we measured the temperature at the amplifier-MEA interface using the TC02 (Multi Channel Systems, Reutlingen, Germany) and adjusted the incubator set point accordingly. The incubator was typically set to 34.5 °C to compensate for heating by the recording devices and NeuroDisco, which increased the temperature by approximately 2 °C and 0.6 °C, respectively.

Electrical activity on CMOS-MEAs was recorded using the MaxOne System (MaxWell Biosystems, Zurich, Switzerland). The MaxOne HD-MEAs comprise 26,400 electrodes per MEA (17.5 µm pitch, electrode size of 8.75 × 12.50 µm^2^ and 1,024 recording channels). Data were recorded using the MaxLab Live software (Ver. 25.1.8.2; MaxWell Biosystems, Zurich, Switzerland) with a sampling rate of 20 kHz and a default high-pass filter of 300 Hz. Except for the high-density recording shown in Fig. 4f, electrode routing was performed by selecting 1,024 electrodes evenly distributed across the recording area. For the high-density recording, a localized region comprising a 23 × 23 electrode array was selected and routed.

Illumination-induced artifacts on MEA electrodes can be mitigated by standard band-pass filtering even at irradiances of about 15 mW/mm^2^ (Kobayashi et al., 2024); at the roughly 1,000-fold lower irradiances used here, we observed no opto-electrical artifacts in high-pass-filtered recordings. In some recordings with the CMOS-based HD-MEAs (MaxWell Biosystems), we were able to induce artifacts shining 470 nm light of 1 mW/mm^2^ onto the amplifier region of the chip, outside the recording area. We therefore never expose this region during our experiments.

### Patch Clamp recordings

For whole cell patch clamp recordings, neuronal cultures were plated on 18 mm coverslips or CMOS-MEAs. On DIV 21-28, coverslips were transferred to a heated recording chamber in an upright microscope (Axio Examiner, Zeiss, Germany) with a 1-2.5 ml/min flow rate of oxygenated artificial cerebral and spinal fluid (ACSF, in mM: 125 NaCl, 4 KCl, 26 NaHCO3, 10 glucose, 1.3 MgCl2, and 2 CaCl2). All recordings were performed at 36 ± 1 °C. Borosilicate pipettes with a resistance of 4-6 MΩ were pulled using a horizontal DMZ-Zeitz-Puller (Zeitz, Germany) and filled with internal solution (in mM: 135 K-gluconate, 10 KCl, 4 NaCl, 0.1 Na4EGTA, 1 Mg-ATP, 0.3 Na-GTP, 10 HEPES, 0.5 Na2-phosphocreatine, and 0.3% (wt/vol) biocytin; 285 to 290 mOsm, pH adjusted to 7.3). Whole-cell current clamp recordings were made using an EPC10 USB amplifier controlled by Patchmaster software (HEKA Elektronik). Fast and slow capacitances were carefully adjusted. Series resistance was 90-100% compensated with a feedback time constant of 10 µs. Voltage signals were low-pass filtered at 8.8 kHz and digitized at 100 kHz. Individual burst events were recorded during 2-10 min gap-free recordings at resting membrane potential. Analysis was performed in the software Igor Pro 9 and 10 (Wavemetrics). Voltages are not corrected for liquid junction potential. Correction would shift them towards more negative potentials by 14 mV.

### Stimulation protocols

#### Stimulus creation and presentation in the flexND

The flexND presented a new pattern every 62.5 ms (16 frames/s). In each frame, the on/off state of each of the 256 LEDs was drawn independently, with an on-probability of 8% or 25%; this gives spatiotemporally uncorrelated Poisson-Bernoulli patterns. The random process was initialized with a fixed seed, so that the same seed reproduced the identical pattern sequence (frozen noise). Global LED light power was set with a control voltage of between 5.2 and 6.0 V (Supplementary Fig. 1). On-probability and brightness were held constant within an epoch. During dark epochs, all LEDs were off.

#### Alternating stimulation protocol (Figs. 2, 3, 4a,e and 5e,f)

Each recording lasted 1,075 min. After 60 min of spontaneous activity, 96 illumination epochs of 270 s (4,320 frames each) alternated with 95 dark epochs of 270 s (about 860 min in total), followed by about 155 min of spontaneous activity. The six stimulation conditions, combining two spatial coverages (on average 8% or 25% of the LEDs switched on per frame) with three LED controller voltages (5.2 V, 5.5 V and 6 V) were each presented 16 times: four frozen pattern sequences (A-D, defined by their seeds) were each shown four times per condition. Note that the same pattern sequences were shown at all three light power values. The order of the 96 illumination epochs was randomized to avoid sequence effects. We obtained 28 such recordings from cultures around DIV 21.

#### Frozen-noise protocol with four light levels (Fig. 5a-d)

In a second set of 20 recordings, 1 h of spontaneous activity was followed by a stimulation period of 912 min (15.2 h) and a further hour of spontaneous activity. The stimulation period comprised 192 illumination epochs of 135 s, each followed by a 150 s dark epoch. The on-probability was 8% in all epochs. Each epoch presented one of two frozen-noise sequences (A or B) at one of four LED controller voltages (5.4, 5.6, 5.8, or 6.0 V). Sequences and voltages were presented in randomized order, so that each of the eight sequence-voltage combinations was repeated 24 times.

#### Long-term viability experiment (Supplementary Fig. 3)

Cultures were randomly assigned to a chronic-stimulation group or a control group. Both groups were prepared from the same litters and expressed ChR2. On DIV 14 and 15, both groups were recorded and otherwise treated identically. After the recording on DIV 15, the chronic-stimulation group was stimulated continuously with the flexND until the recording on DIV 22 (7 days). The stimulus was the uncorrelated Poisson-Bernoulli patterns described above (62.5 ms frames, on-probability 25%, control voltage 5.5 V). It ran without dark epochs. Stimulation was interrupted only for medium exchange (DIV 14, 17 and 21) and for the spontaneous part of each recording. Stimulated cultures remained on their devices throughout; the temperature measured at the MEA was 36-37 °C. Control cultures were kept in a separate incubator at 36-37 °C. For each recording, they were transferred to the incubator of the stimulated cultures and recorded with the same flexND and MEA2100-Mini setup. After the recording on DIV 22, chronic stimulation was stopped and the group was treated like the controls. Recordings were made on DIV 14, 15, 16, 17, 18, 22, 23 and 24. Control recordings lasted 45 min: 15 min of spontaneous activity, 15 min of stimulation with the same parameters, and another 15 min of spontaneous activity. While chronic stimulation was ongoing (DIV 16-22), the stimulated group was recorded for 30 min: the last 15 min of ongoing stimulation followed by 15 min of spontaneous activity. On all other days, both groups were recorded with the 45 min protocol. Each full recording was spike-sorted and every unit separated as its own cluster was counted as active. Because control recordings were longer, this comparison, if anything, favors higher unit counts in the control group. For each DIV, the number of active units was compared between groups with two-sided Mann-Whitney U tests, Holm-corrected across the eight DIVs (12-24 cultures per group and DIV).

#### Correlated stimuli (Fig. 4e)

In a small set of experiments, correlation was introduced to the stimuli in the following way. Each pattern (or frame) is drawn as described before with an on-probability of 8%. Next, the on-state of a set of two master-LEDs is monitored. If a master LED is lit in the current pattern, the on-state of a set of 8 slave LEDs is redrawn, this time with an on probability of 80%. The eight slave LEDs and their master LED form either a straight line, or a 3 × 3 block. In these experiments, 16 h of stimulation with partially correlated patterns were bracketed by more than 1 h of stimulation with fully uncorrelated patterns at 8% on-probability. Throughout a given experiment, the LED controller voltage was held constant: 5.8 V in 5 experiments, 6.0V in two others.

#### Stimulus creation and presentation in the HD-ND

With the current setup, all stimuli are created in real time. The stimuli can be created in two different ways: Python code can create frame contents of arbitrary complexity. It is stored in memory and from there routed to the GPU with a C package. The GPU shader can create simpler frame content that can be described by parametric patterns, e.g., sinusoids. They require no extra transfer step and can be created and presented at 60 Hz, while the Python-based route reaches 22.5 Hz for the Poisson stimuli we apply in this study.

#### RF estimation with HD-ND (Fig. 4b-d,f)

The reverse correlation experiments used Bernoulli-Poisson patterns with block sizes of 92, 47.5, and 19 µm. The on-probability was always 10%, and the brightness was maximal (255). For the highly resolved RFs (Fig. 4c,d), stimuli with block sizes of 19 and 47.5 µm were presented for 1 h. All experiments were performed on cell cultures at DIV 20-21.

#### Drifting sinusoidal gratings

To assess orientation selectivity, we recorded activity from 1,024 electrodes in a single culture at DIV 20 during presentation of drifting sinusoidal gratings. We rendered the gratings with a custom OpenGL fragment shader using the following equation:

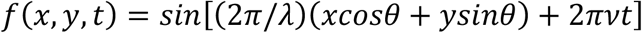

Here, x and y are coordinates in the culture plane, t is time, and θ is the angle of the spatial wave vector. The spatial wavelength was λ = 760 µm. The temporal frequency was ν = 2.0 Hz.

We presented ten epochs, each with gratings of 16 directions (θ = 0° to 337.5° in 22.5° increments). Each grating was presented for 4 s, followed by 3 s of darkness. Epochs were separated by an additional 2 s of darkness. Because the firing rate was transiently increased throughout most of the first epoch, it was excluded from the analysis.

### Data Analysis

The procedures detailed in the following section are realized by software modules available on :::::. The granular details are explained in inline comments inside that code.

#### Spike Detection and Spike Sorting

For data acquired on 60-electrode MEAs spike sorting was performed with a custom pipeline implemented in Python, using Dask for parallel computation and Zarr as the storage backend for parallel I/O. This framework enables continuous analysis of the entire 18 h recordings (sampled at 50 kHz, 300 GB) as larger-than-memory datasets. The same code runs on laboratory laptops and HPC systems, with resource allocation matched to data volume and identical results across platforms. The continuous signals were preprocessed (reference subtraction, whitening) and filtered with a configurable, acausal filter bank combining SciPy-based Butterworth band-pass with notch and comb filters (e.g., against 50 Hz line noise and its harmonics), whose joint frequency response can be inspected directly.

In the preprocessed data, spike snippets were defined as crossing the threshold of 4.5 standard deviations. Spike snippets were clustered per channel using k-means on a combined feature space of principal components of the whitened waveforms, principal components of their frequency spectra, a UMAP embedding, and waveform descriptors (e.g., the deflection from threshold crossing to peak). For each channel, the optimal cluster number k was determined by consensus across complementary cluster-quality metrics (silhouette score, Calinski-Harabasz index, Davies-Bouldin index), identifying k candidate units. Finally, an automated putative-unit classification (unit vs. noise) was applied to these clusters and validated against manual curation.

For data acquired on the CMOS-MEAs, the spike threshold was set to 5 times the RMS of the noise level. The raw recording data were subsequently analyzed to identify the corresponding electrode IDs using the open source Python package SpikeInterface (Buccino et al., 2020).

#### Instantaneous firing-rate estimation (Fig. 2)

Population instantaneous firing rates were estimated from the pooled spike trains of all detected units using Neo and Elephant ^54^. Spike times from all units were concatenated, sorted, and represented as a Neo SpikeTrain. The pooled spike train was convolved using Elephant’s instantaneous_rate function with a Gaussian kernel. The Gaussian standard deviation and temporal sampling period were both set to 1 s, close to the standard deviation estimated with ^55^. The resulting population rate was divided by the total number of recorded units, yielding the mean instantaneous firing rate per unit.

#### Selection of intervals for burst detection and MFR calculation

Burst detection and MFR calculation were estimated in the stationary final 160 s of each stimulation epoch. All spontaneous intervals before the first non-spontaneous interval and all spontaneous intervals after the final non-spontaneous interval were included in full. For spontaneous intervals located between stimulation conditions, the final 160 s were analyzed. Detection of the initial burst after each stimulation onset (Fig. 3f) used the same threshold as detection of the other bursts of the same epoch.

#### Mean firing rate estimation (Fig. 2, Supplementary Fig. 5 and Supplementary Table 1)

The per-epoch mean firing rate was calculated with:

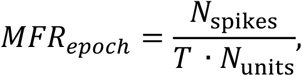

where *N*_spikes_ is the total number of spikes in the epoch, *T* is the epoch duration in which the MFR is calculated, and *N*_units_ is the total number of spike-sorted units.

#### Relative Firing Rate Estimation (Fig. 2b)

For each neuron, the firing rate in each 54 s time bin was normalized to its mean firing rate during the first hour of recording.

#### Population-burst detection

Population bursts were detected using a two-stage procedure based on unit-specific burstlets and their temporal coincidence across units, adapted from Wagenaar et al. (2006). For every unit and stimulus interval, a burstlet threshold was calculated from a dedicated inter-spike interval (ISI) estimation window. For unit *i*, the threshold was

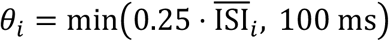

where 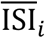 denotes the mean ISI within the corresponding estimation window of that unit. The resulting threshold was subsequently applied to the spikes in the corresponding burst-detection window. A unit-level burstlet was defined as a sequence of at least four spikes for which all consecutive ISIs were shorter than *x*_*i*_. If burstlet spikes of at least two units appeared inside a 10 ms window, this window was considered part of a population burst. In a second run, population bursts separated by less than 1 s were merged into a single burst. The bursts defined in this way are the basis of a number of statistics: burst frequency, fraction of all spikes that occur during bursts, mean burst duration, mean number of spikes per burst, intra-burst MFR, and mean number of participating units per burst. A unit was classified as participating if it emitted at least one spike within the burst boundaries.

#### Burst latency ranks and reference construction

Here, we aim to characterize the sequence with which individual units were recruited to the initial progression of the burst. The first step identifies all spikes that are relevant to this process. We decided to include all spikes fired within a 100 ms window starting 5 ms before the detected burst onset.

For every unit *i* and detection window *b* = [*t*_0,*b*_, *t*_1,*b*_), first-spike latency *L*_*i,b*_ was defined as

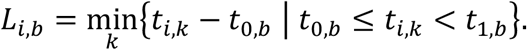

The first-spike latencies *L*_*i,b*_ of all participating units were converted to zero-based ranks *R*_*i,b*_ with earlier spikes receiving lower ranks. Equal latencies were assigned their average rank (midrank); numerical equality was evaluated using a tolerance of 10^−1*2*^ s, a value between the float uncertainty and sampling timestep.

We can compare the temporal progression of two bursts based on their rank sequences. For a global analysis across all bursts in a recording, a common reference sequence is needed (Fig. 3d-f). This consensus reference was constructed from all bursts detected in the first 276 s of spontaneous activity of the recording. Because the number of participating units could differ between these reference bursts, ranks were normalized separately within each reference burst:

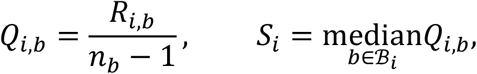

where *R*_*i,b*_ is the zero-based rank of unit *i* in reference burst *b, n*_*b*_ is the number of ranked units in that burst, and *ℬ*_*i*_ is the set of reference bursts in which unit *i* participated. The finite consensus scores *S*_*i*_ were converted to midranks to obtain the final reference vector.

#### Burst-rank, aka intra-burst trajectory correlations (Fig. 3)

Similarity between two bursts was calculated using only units that appear in the rank sequences of both bursts. Burst pairs with fewer than five overlapping units were assigned no correlation value.

Kendall’s tau-b was used as the primary rank-correlation coefficient. Tau-b explicitly accounts for ties in either rank vector. Thus, equal first-spike latencies and equal reference ranks were retained as ties rather than resolved arbitrarily.

#### Hierarchical clustering of burst-similarity matrices (Fig. 3g,h)

All-to-all pairwise burst-similarity matrices were ordered by agglomerative hierarchical clustering. Pairwise correlation values *C*_*ij*_ were transformed into dissimilarities according to

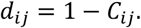

Consequently, perfectly correlated bursts had a dissimilarity of 0, uncorrelated bursts a dissimilarity of 1, and perfectly anticorrelated bursts a dissimilarity of 2. Correlations that were undefined because of insufficient overlap or constant ranks were assigned the maximum dissimilarity of 2. Hierarchical clustering was performed using average linkage as implemented in SciPy. The distance between two clusters was therefore defined as the average of the pairwise dissimilarities between their members. Optimal leaf ordering was enabled to minimize the summed dissimilarity between adjacent leaves while preserving the hierarchical cluster tree.

#### Receptive Fields

Let the stimulus vector at time *t* be denoted as *x*_{*t*}_, containing all spatial pixels and the temporal history in a fixed window. The *raw stimulus ensemble* is the distribution P(x_{t}_) across all times and the *spike-triggered ensemble P*(*x*_{*t*}_ |spike) is the distribution of vectors in a fixed window around each spike.

The spike-triggered average (STA) stimulus is the simplest estimate of the stimulus filter. The idea is to identify deviations between the spike-triggered ensemble *P*(*x*_{*t*}_ |spike) and the raw stimulus ensemble *P*(*x*_{*t*}_), thereby revealing those stimulus features that influence spiking. The STA captures the first moment of the spike-triggered ensemble by comparing it to the raw stimulus ensemble. Here, *x*_{*t*}_ is the stimulus vector at time *t*, containing all spatial pixels and the temporal history. Let *t*_{*n*}_ denote the time of the *n*-th spike. An STA recovers one linear filter (with non-zero mean projection) and following Schwartz et al. (Schwartz 2006) is given by

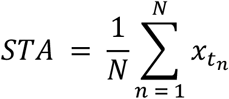

#### Normalization

Each pixel of the STA was expressed as a z-score against the raw stimulus ensemble,

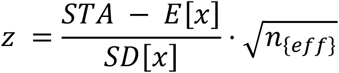

*E*[*x*] and *SS*[*x*] are the mean and standard deviation of that pixel over the frames in the analysis. Here *n*_*eff*_ is not the spike count *N* of the equation above, because the sub-frame bins of one frame would repeat the same pattern. Spikes falling in the same frame are not independent draws from *P*(*x*_*t*_|*spike*). Instead, *n*_*eff*_ is the effective number of stimulus presentations behind the average (Kish’s effective sample size). Let *m*_*i*_ be the number of spikes the unit fired during frame *j, J* the number of frames entering the analysis, *N* = ∑_*i*_ *m*_*i*_ and 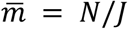. Then

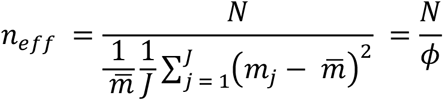

Here 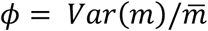 is the Fano factor of the spike count per frame. A unit that rarely fires more than one spike per frame has *ϕ* ≈ 1 and *n*_*eff*_ ≈ *N*. A bursting unit delivers several spikes to the same picture, has *ϕ* > 1 and therefore *n*_*eff*_ < *N*. The mean 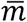 is subtracted because *E*[*x*] is estimated from the same frames; hence, only the variation of spike count about its mean adds uncertainty. A z-score grows with 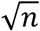. Quantitative comparisons between two maps obtained for different numbers of spikes are therefore not based on the z-scores, but on the effect sizes (Supplementary Fig. 8):

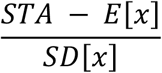

#### Window and binning

For all stimuli presented with the flexND, one stimulus frame lasted 62.5 ms. For the analysis, each frame was divided into six bins of 10.42 ms, each carrying the pattern of its own frame, so that a bin never reaches across a frame boundary. The STA analysis window of *x*_*t*_ ran from ten bins before to five bins after the spike (-104 ms to +52 ms relative to the spike). In figures, where only the STA for one single lag value is shown, it always is the bin before the spike, i.e., lag -10.42 ms. Frames without lit LEDS were excluded, as were units with fewer than ten spikes inside the window.

For the HD-ND, the window size was chosen to be identical to the frame update time step because our focus lay on the spatial resolution.

#### Subsampling of spikes (Supplementary Fig. 8a)

The spikes of one unit, recorded within a single 14.7 min Poisson block, were expanded into a list of individual spikes, of which a fraction *f* was drawn without replacement, keeping a subset *N*_*f*_ = ⌊*f* · *N*⌋ of the spikes. The retained spikes were binned again, and the map recomputed exactly as for the full set. Each subset was drawn sequentially without replacement; they share no spikes.

#### Biphasic profiles

A unit was counted as biphasic when the z-value at the strongest pixel of its map changed sign over the STA analysis window.

#### Construction of extracellular footprint (Fig. 4f and Supplementary Fig. 9)

The extracellular signal of the CMOS reflects neuronal activity in the immediate surrounding. In analogy to the calculation of the STA stimulus, we determine an STA of the extracellular signal.

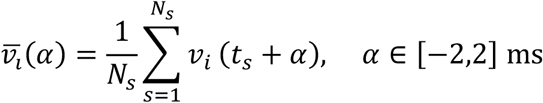

where v_i_(t) is the voltage recorded at electrode i, t_s_ is the time of the s-th spike, N_s_ is the total number of spikes, and α is the time relative to the spike time. A symmetric window of ±2 ms around each spike was used to capture the extreme voltages, also at locations where the maximal activity was slightly earlier or later than at the soma, the presumed triggering location. We construct the footprint from the maximal voltage values of 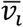 of all electrodes. This footprint captures all electrical activity that is tightly correlated to the spiking of that neuron, in particular the depolarization of its dendrites and axon. Because the amplitudes of dendritic, somatic, and axonal signals vary by at least an order of magnitude, we use a logarithmic color scale.

## Ethics Approval

All procedures were performed in accordance with German law on animal research and registered with the animal protection offices of the University Medical Center Göttingen and the University of Göttingen under the licenses T 23.13 and T2-2025.

## Data Availability

Raw data for all figures will be stored in a public repository before submission. flexND PCB plans and 3D printer files for the Lid and HDND body are given in the supplemental documents.

## Code Availability

Code will be available upon submission at github.com/goecidbn/

## AI Usage

Large language models were used for literature search and readability improvements.

## Acknowledgement

We thank Tristan Desgardin for initial analysis of firing statistics and synchronization issues. We thank Yonathan Katz for inspiring discussions on optogenetics in cultures and an initial design of the lid. We thank Johannes Zierenberg and Darius Burr for valuable input on analysis and simulation. We thank Marion Kunze, and Sven Franz for workshop assistance and Heidi Brodmerkel for technical assistance with the cell cultures.

We thank Dr. Kathrin Kusch (Institute for Auditory Neuroscience, Göttingen) and the Viral Core Facility (VCF) of Charité - Universitätsmedizin Berlin (RRID:SCR_027965) for providing the viral preparations for ChReef and ChR2, respectively.

CSA and FS were supported by the Else-Kröner-Fresenius Doctoral College of the University Medical Center Göttingen. CSA, FS and HZK were supported by the German Academic Scholarship Foundation (Studienstiftung des Deutschen Volkes).

This work was supported by the Bundesministerium für Bildung und Forschung (BMBF, Federal Ministry of Education and Research) grant 01GQ0811, 01GQ01005B, 01GQ0922; Volkswagen Foundation ZN2632 & “Niedersächsisches Vorab”. Deutsche Forschungsgemeinschaft (DFG, German Research Foundation https://www.dfg.de/en) under Germany’s Excellence Strategy - EXC 2067/1- 390729940 (to FW), and Project ID 528760423 (CRC 1690) (to AN and FW).

## Notes

### Competing Interest Statement

The authors have declared no competing interest.

