## Extended Data for "NeuroDisco: Interfacing neuronal populations - cell by cell, millisecond by millisecond, day by day"

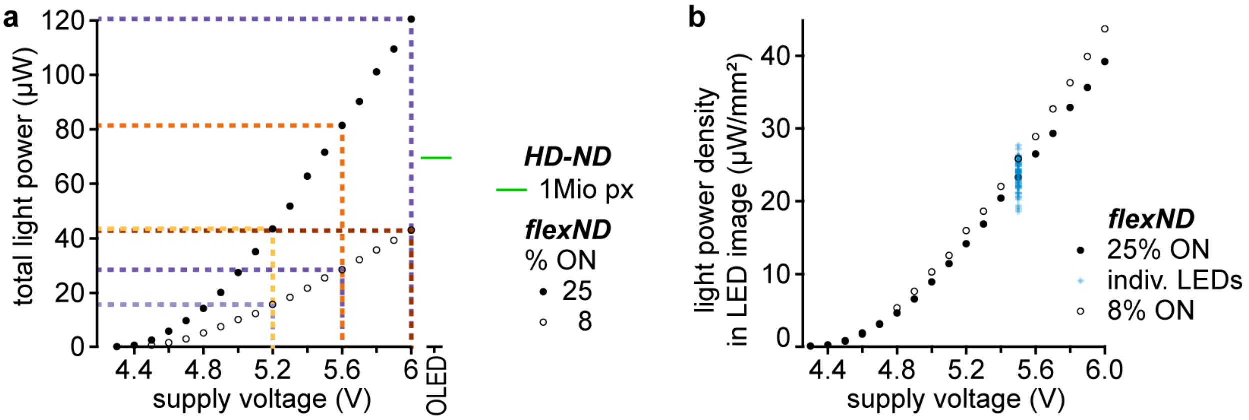


**Supplementary Figure 1 | Total power and approximate light power density of NeuroDisco stimuli.** **a,** The light power in the entire image with the specified configuration, determined with a calibrated photodiode (S2938-K, Hamamatsu; see Methods). Colored dashed lines correspond to the intensity-coding colors in Fig. 2a. Orange shades refer to 25% coverage, purple to 8%. **b,** The light power densities approximated from images with the Raspberry Pi HQ camera V1.0. The numbers presented here are for fully focused images (see Supplementary Fig. 4) and under the assumption of linear light sensitivity of the camera.

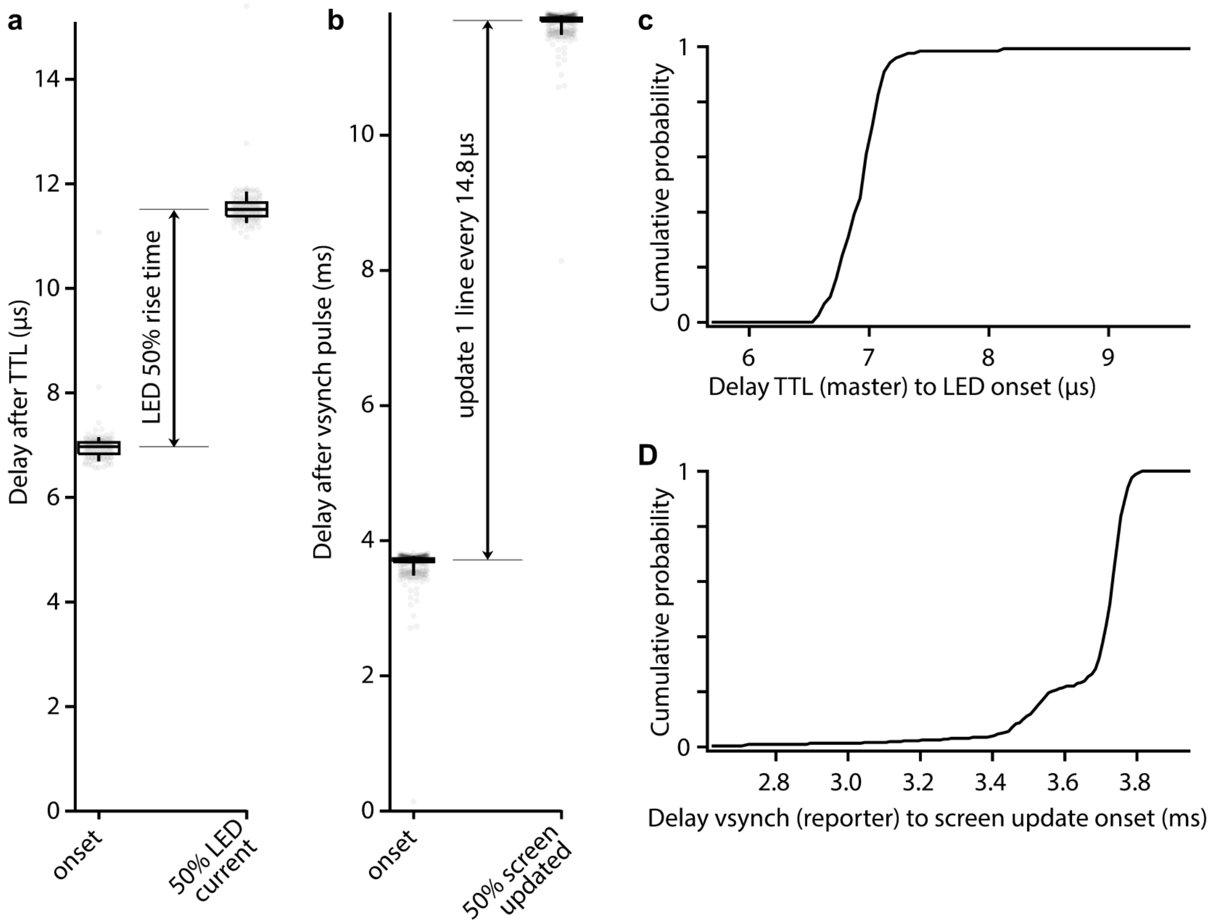


**Supplementary Figure 2 | Synchronization of ND output. a,** Delay between TTL "Next frame" command to Arduino pin and light onset (left column), respectively 50% light output (right column). **b,** Delay between the vsync signal of the Raspberry Pi Graphics Processing Unit and the screen update onset (left), respectively the update of 50% of the screen (right). Updating all 1,080 rows takes 16 ms. a and b show individual data points and Tukey Box plots with boxes encompassing the 9th to 91st quantile. **c,** Cumulative data of the left row in a. Median delay to onset is 6.97 µs with interquartile range 0.26 µs. **d,** Cumulative data of the left row in b. Median delay to onset is 3.73 ms with interquartile range 0.84 ms.

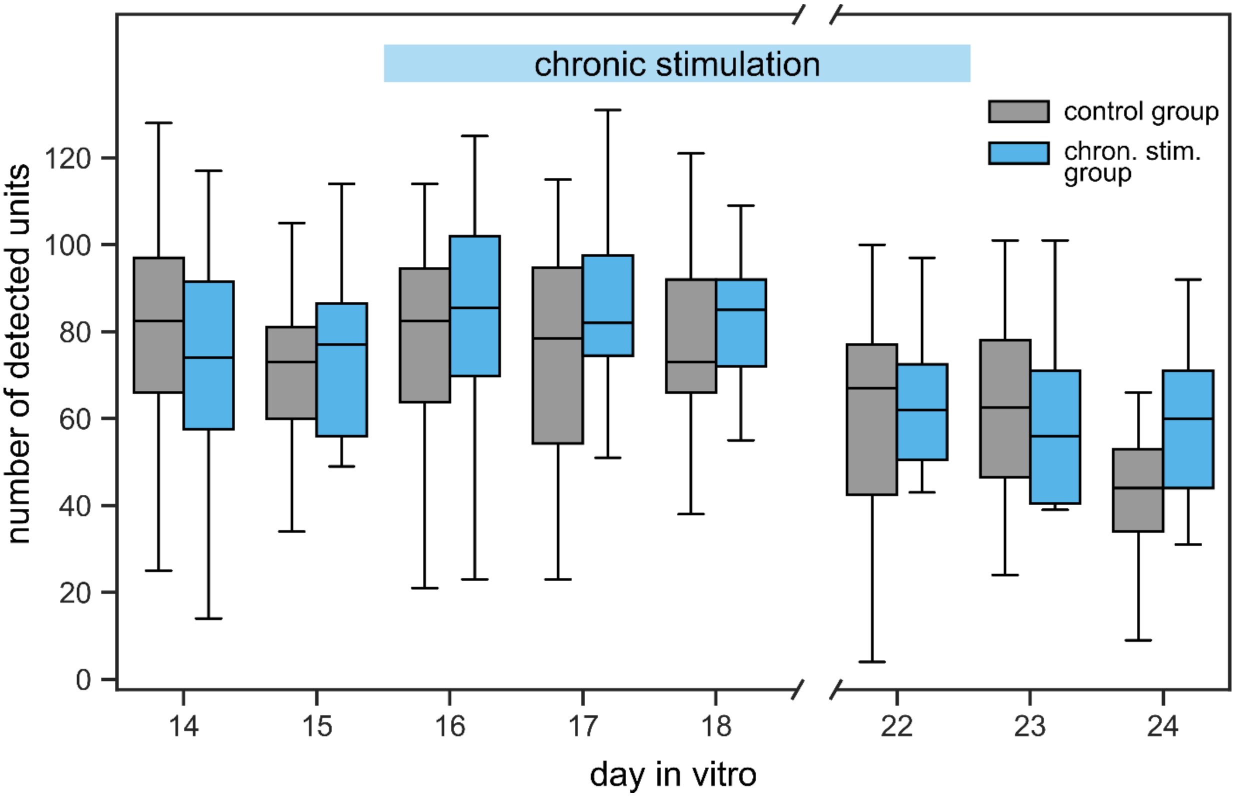


**Supplementary Figure 3 | Chronic stimulation does not reduce yield of detected neurons after spike sorting.** Number of spike-sorted units obtained from two groups of cultures on MEAs i) control group, only stimulated for 15 min per day during electrical recording and ii) stimulation group, chronically stimulated starting after the measurement at 15 days in vitro (DIV 15) until after the measurement at DIV 22. Stimulus was coverage 25%, 5.5 V voltage at LED drivers. The light-blue bar at the top denotes the interval of 7 days (DIV 15.5–22) during which the stimulation was applied continuously for the chron. stimulation group, except for a 15 min interval per measurement day and medium exchanges that happened on DIV 14, 17 and 21 after recording. Data are shown as Tukey boxplots. Multiple observations from the same identifier at the same DIV were averaged before statistical analysis. Control and chronic-stimulation groups were compared separately at each DIV using two-sided Mann–Whitney U tests, with Holm correction across the eight comparisons. No significant group differences were detected (all Holm-adjusted $P = 1.00$; minimum unadjusted $P = 0.189$); sample sizes ranged from 12 to 24 cultures per group and DIV.

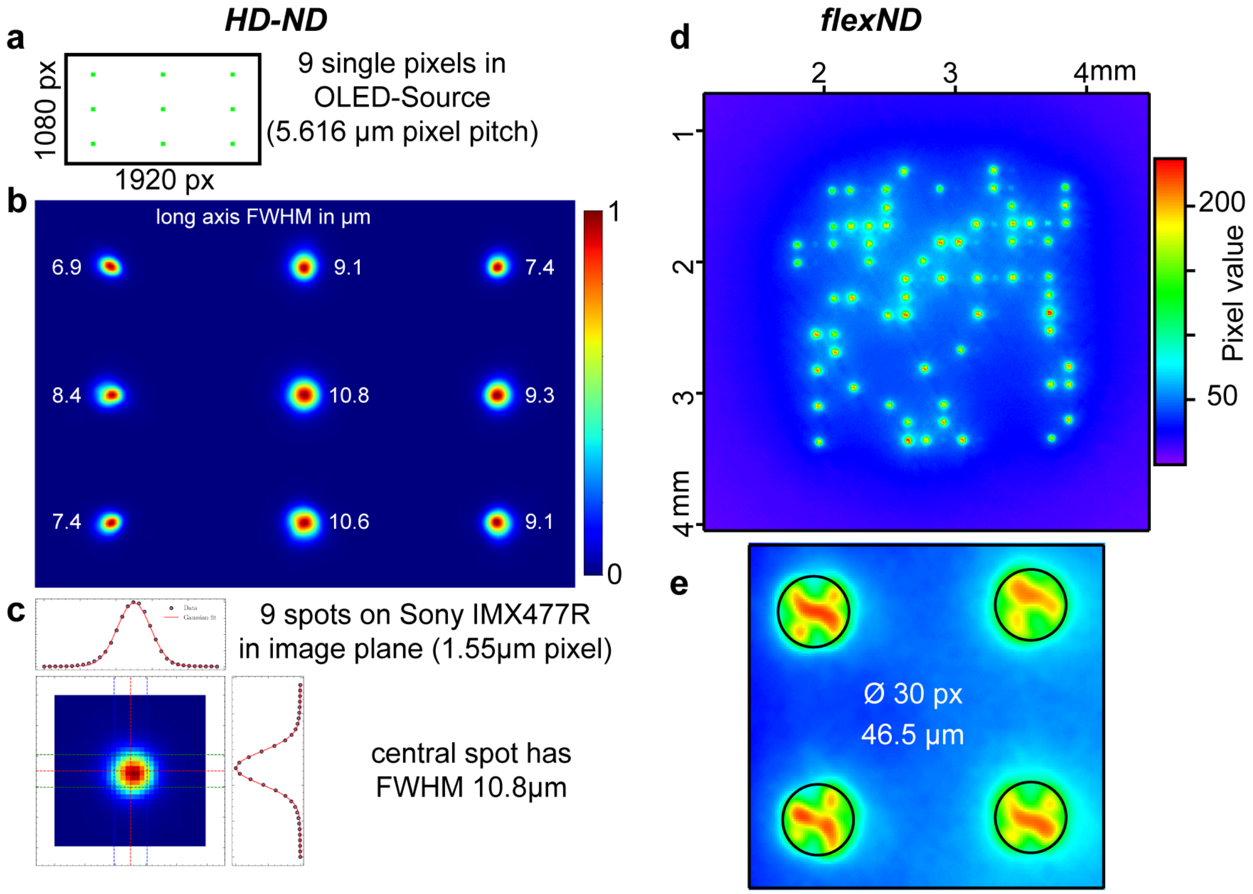


**Supplementary Figure 4 | The spatial structure of optical stimuli. a,** The nine-point matrix used to probe the HD-ND's point spread function. **b,** The image of a captured with the Raspberry Pi HQ camera V1.0. White numbers indicate the full-width at half-maximum along the longer axis of the nine spots. **c,** Gaussian fits of the intensity profiles of the central spot. **d,** One frame with 25% coverage (64 LEDs ON) taken with the Raspberry Pi HQ camera V1.0 image sensor at the focal plane of the flexND. **e,** A zoom into a similar picture reveals the fine structure of the µLED emission pattern. The black circles have a diameter of 46.5 µm.

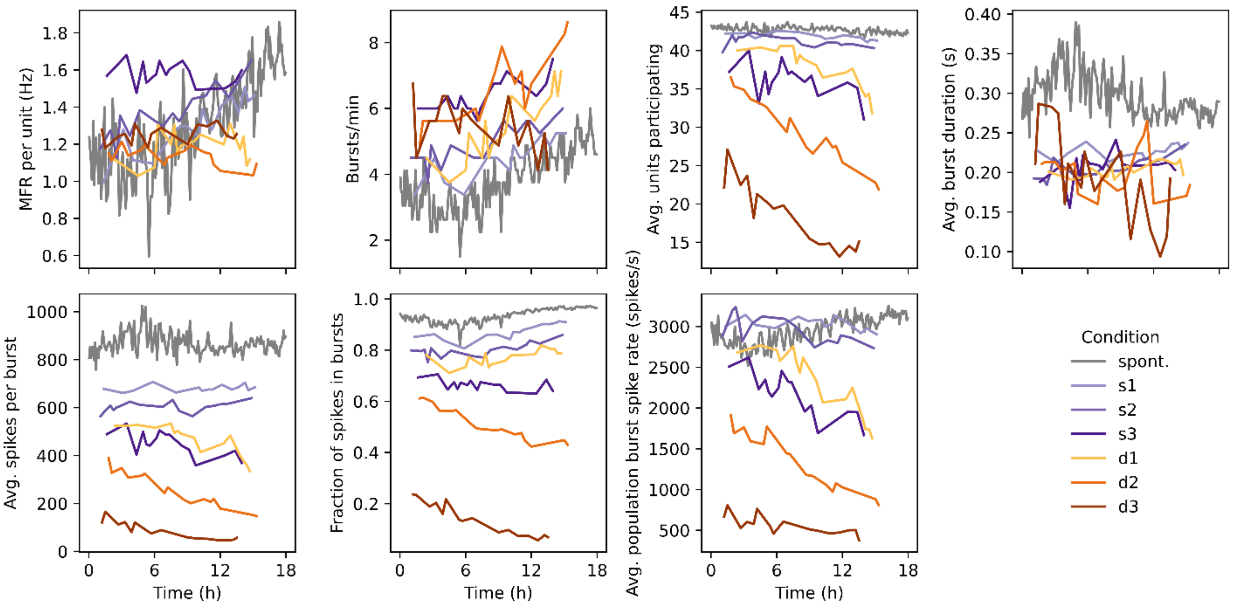


**Supplementary Figure 5 | Burst metrics during various stimulation conditions across 18 h.** Same recording as in Fig. 2. Cultures were recorded for 18 h, 3.5 h without stimulation (spontaneous; sp), 14.3 h alternating darkness and light-stimulation with the flexND. Stimulation conditions varied in light intensity and spatial coverage (8%, s1–s3, purple shades; 25%, d1–d3, orange shades, same color code as in Fig. 2a). Network activity metrics over the 18 h recording, calculated separately for the stationary part (final 160 s) of each 270 s epoch. Data points are connected by straight lines.

| Activity | Stimulus coverage (%) | Voltage (V) | Mean firing rate (Hz) |
|---|---|---|---|
| Spontaneous | – | – | 1.322 |
| Light-driven | 8 | 5.2 | 1.307 |
| Light-driven | 8 | 5.5 | 1.364 |
| Light-driven | 8 | 6 | 1.576 |
| Light-driven | 25 | 5.2 | 1.185 |
| Light-driven | 25 | 5.5 | 1.133 |
| Light-driven | 25 | 6 | 1.253 |

**Supplementary Table 1 | Mean population firing rates during spontaneous and light-driven activity.** The grand mean of population firing rates was calculated across repeated presentations. Firing rates during the period of alternating stimulation were calculated excluding the onset response (only the last 160 s of each block were included). The spontaneous reference was calculated across analysis windows spanning pre-stimulation, inter-block, and post-stimulation activity.

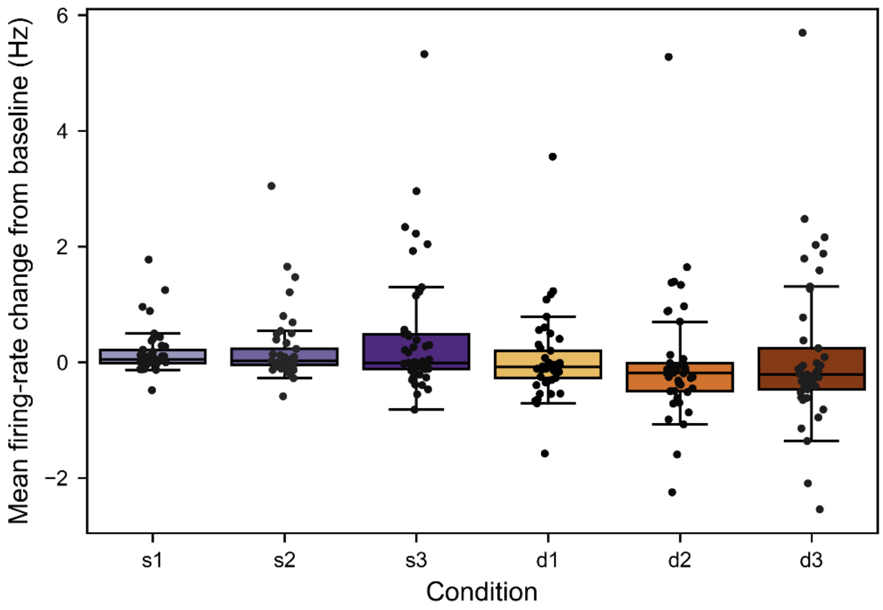


**Supplementary Figure 6 | Single neuron firing-rate changes relative to spontaneous baseline across stimulation conditions.** For each unit, firing rate was calculated from spike counts during the final 160 s of each stimulation period and averaged across presentations of the same pattern–brightness condition. The unit's mean firing rate during the initial spontaneous baseline was then subtracted to obtain the change in firing rate (Hz). Data shown as Tukey boxplot.

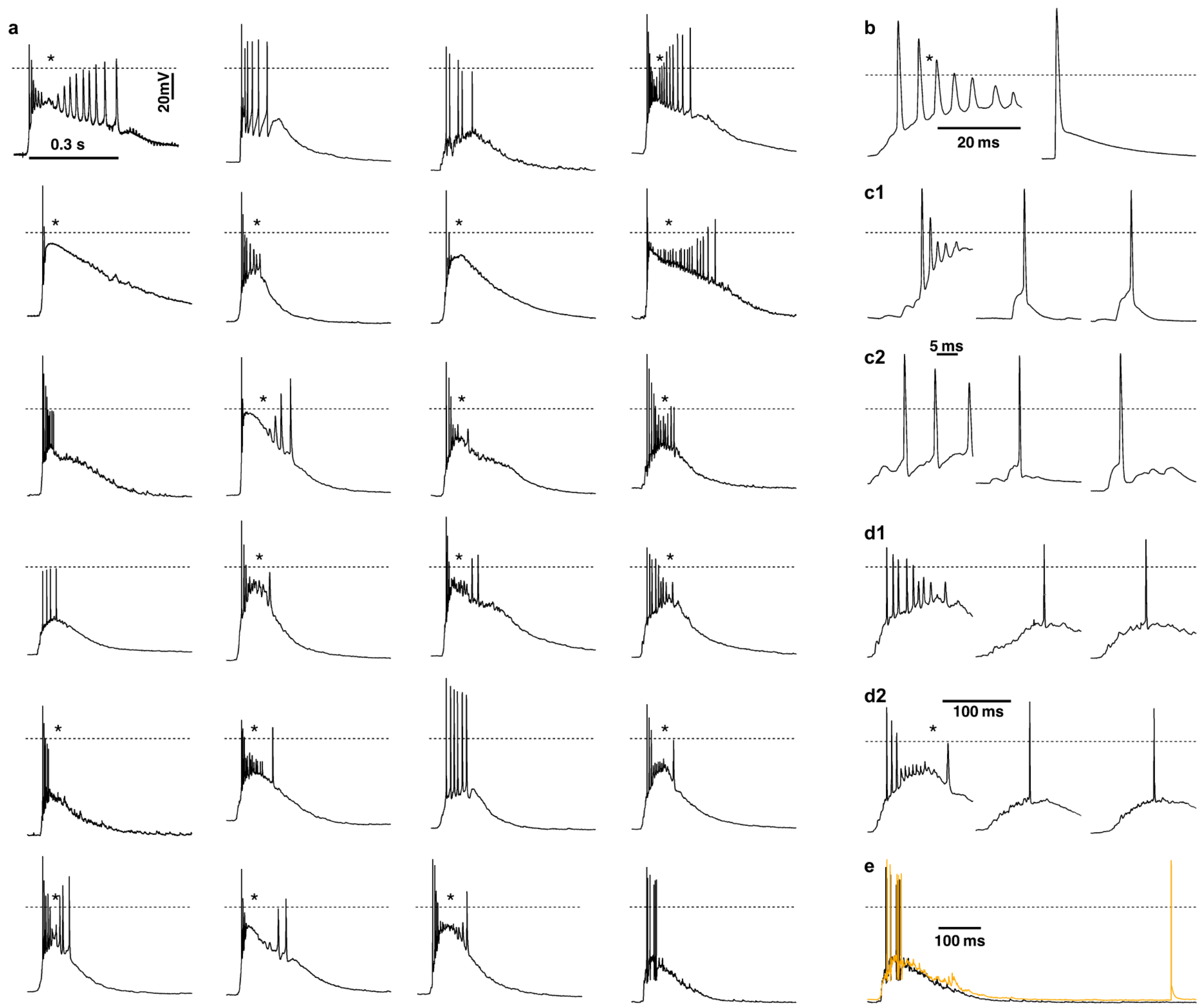


**Supplementary Figure 7 | Population bursts in neuronal cultures drive most neurons into depolarization block (paroxysmal depolarizations). a,** Whole-cell current-clamp recordings from 24 neurons in 24 neuronal cultures from 3 preparations. Each trace is representative of the responses we recorded from this particular cell. The responses on the top left and the one just below are recorded during simultaneous extracellular recordings with 1,000 channels on a HD-MEA. For these we have independent confirmation that population-wide bursts cause these depolarizations. For the other recordings, this is the most parsimonious explanation. The asterisks denote traces that show clear signs of depolarization block, e.g., strongly reduced AP amplitude, increased AP width, silent periods during strong depolarization, preceded or bracketed by activity during periods with less depolarization. **b,** Magnification of the burst response just to the left of it in a, and a single action potential from the same cell, fired outside a burst. Note how the action potential width and height change during the progress of the burst and how narrow the single action potential is. **c1,c2,** Magnification of the onset of the bursts depicted just left of the traces in a and, for each cell, two action potentials fired apparently in response to a single excitatory input, outside bursts. **d1,d2,** Magnification of the bursts depicted just left of the traces in a and, for each cell, two responses to more rare, smaller bursts that did not cause depolarization block. **e,** Overlay of two bursts recorded in the same cell. Note the similarity of the overall depolarization envelope and the firing pattern. Also note the individual action potential in the orange trace.

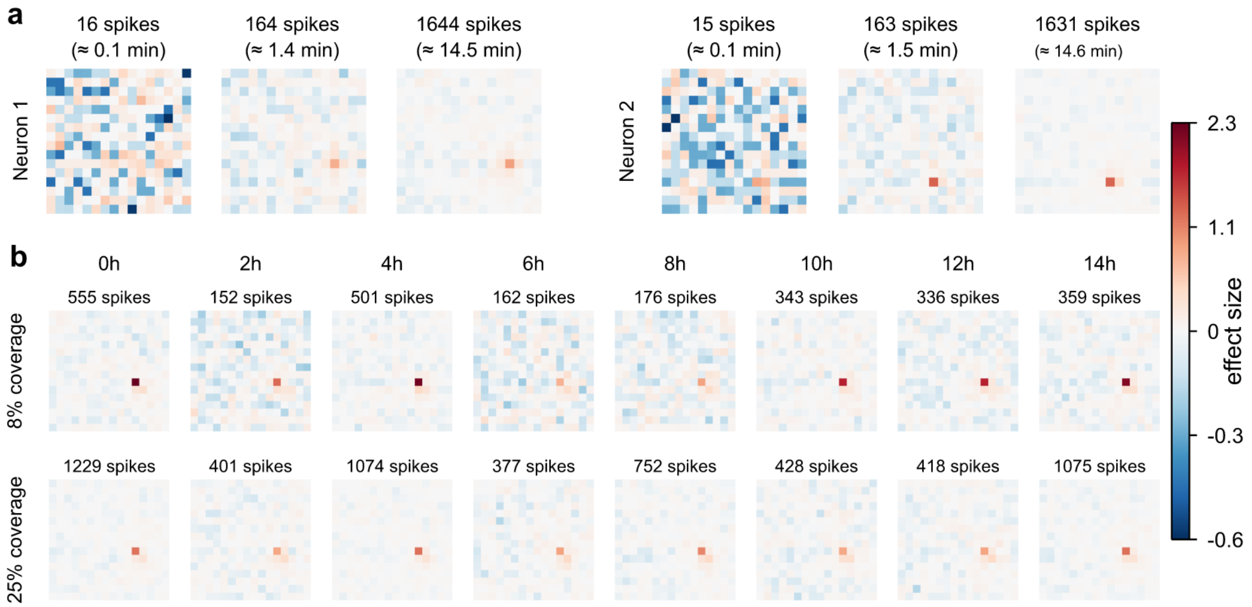


**Supplementary Figure 8 | Receptive-field stability and spike count dependent qualtiy. a,** To illustrate how RF quality depends on spike count, RFs were reconstructed from randomly subsampled spike numbers for two representative neurons (1%, 10% and 100% of the spikes in one 14.7 min Poisson block). The neurons fired at 1.85 Hz, so the three RFs correspond to intervals of 9, 88, and 880 s. Color is the effect size: the deviation of the spike-triggered average from the mean stimulus, in units of the standard deviation of the stimulus at that pixel. Unlike the otherwise used z-score, it does not grow with the number of spikes. (Methods) **b,** Long-term RF reconstruction across 14.3 h of continuous recording of the type introduced in Fig. 2. The RFs were reconstructed from 4.3 min stimulation epochs throughout the experiment under both 8% and 25% Poisson stimulation. Each map is calculated from a single stimulus epoch and thus a single intensity. Shown are RFs at 10.42 ms before the action potential time. Color is the effect size as in a.

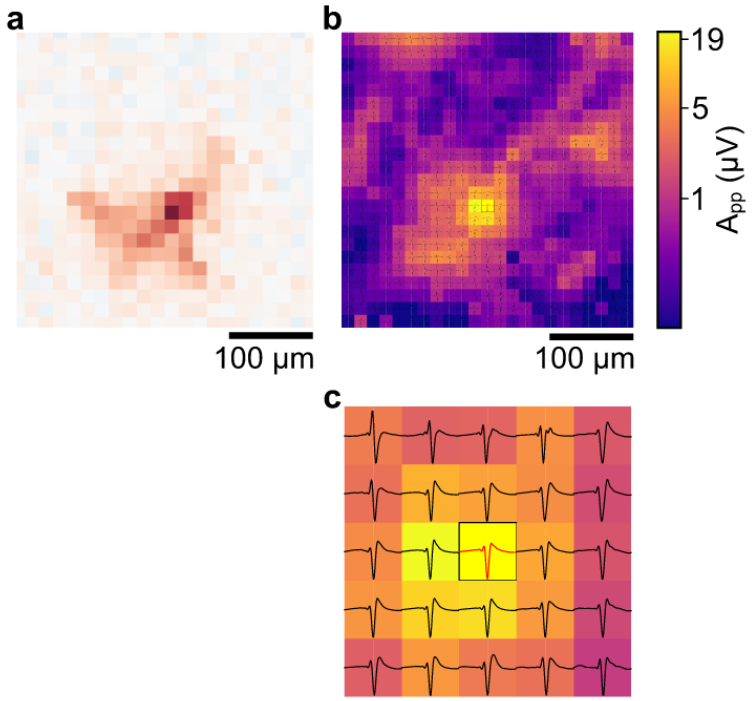


**Supplementary Figure 9 | Supplementary Figure 9 | Spike-triggered extracellular footprint of the neuron in Fig. 4f. a,b,** Data from Fig. 4f shown at higher magnification. Note that each pixel of the electrical footprint contains the normalized waveform recorded at that location, triggered on the peak of action potentials detected at the yellow pixel with the red trace, the triggering unit. **c,** Zoom into 25 electrodes, centered on the triggering electrode.
